# Light activates *psbA* translation in plants by relieving inhibition of translation factor HCF173 by the *psbA* ORF in *cis*

**DOI:** 10.1101/2024.10.26.620444

**Authors:** Rosalind Williams-Carrier, Prakitchai Chotewutmontri, Sarah Perkel, Margarita Rojas, Susan Belcher, Alice Barkan

**Affiliations:** Institute of Molecular Biology, University of Oregon, Eugene, OR 97405

**Keywords:** translational control, PSII repair, *psbA*, chloroplast, HCF136, RBD1, HCF244, HCF173, Arabidopsis

## Abstract

The D1 reaction center protein of photosystem II is subject to photooxidative damage. Photodamaged D1 must be replaced with newly synthesized D1 to maintain photosynthesis. In plant chloroplasts, D1 synthesis is coupled to D1 photodamage via regulated translation initiation on *psbA* mRNA, which encodes D1. Mechanisms underlying this coupling are unclear. We show by analysis of reporter constructs in tobacco that the *psbA* translational activators HCF173 and HCF244 activate via *cis*-elements in the *psbA* UTRs and that the 5’ UTR sequence bound by HCF173 is essential for *psbA* expression. However, the *psbA* UTRs are not sufficient to program light-regulated translation. Instead, the *psbA* open reading frame acts in *cis* to repress translation initiation, and D1 photodamage relieves this repression. A truncated HCF173 mutant conditions constitutively high *psbA* ribosome occupancy in the dark, implicating HCF173 as a mediator of the repressive signal. We propose a model that is informed by structures of the Complex I assembly factor CIA30/NDUFAF1 positing that D1 photodamage relieves a repressive cotranslational interaction between nascent D1 and HCF173’s CIA30 domain, and that the D1 assembly factor HCF136 promotes this interaction. These findings elucidate a translational rheostat that maintains photosynthesis in the face of inevitable photosystem II photodamage.

## INTRODUCTION

Oxygenic photosynthesis is centered in photosystems I and II (PSI and PSII), large protein-pigment complexes embedded in the thylakoid membranes of chloroplasts and cyanobacteria. The absorption of light energy by PSI and PSII is, at once, fundamental to their activities and damaging to the photosynthetic apparatus. The primary casualty of photodamage is the D1 protein in the PSII reaction center. In an elaborate repair cycle, damaged PSII is partially disassembled, damaged D1 is removed by proteolysis, new D1 is synthesized, and PSII is reassembled (reviewed in Jarvi et al., 2015; Theis and Schroda, 2016). D1 damage increases in proportion with light intensity (Sundby et al., 1993; Park et al., 1996; Tyystjarvi and Aro, 1996), with synthesis of new D1 increasing with D1 damage up to moderate light levels. However, at higher intensities, D1 synthesis plateaus and photosynthesis is inhibited (Tyystjarvi and Aro, 1996; Nishiyama and Murata, 2014; Theis and Schroda, 2016). Thus, the rate of D1 synthesis following D1 damage is a key determinant of photosynthetic efficiency.

Despite impressive advances in understanding many aspects of the PSII repair cycle, little is known about how D1 synthesis is activated as needed to provide D1 for PSII repair. D1 is encoded by the *psbA* gene, which is found in the chloroplast genome in plants and algae. It has long been recognized that light activates translation of the chloroplast *psbA* mRNA (reviewed in Sun and Zerges, 2015; Zoschke and Bock, 2018). Recently, this phenomenon in plants was shown to result from the combination of two mechanistically distinct effects of light on chloroplast translation (Chotewutmontri and Barkan, 2018, 2020): (i) light-induced D1 damage triggers the recruitment of ribosomes specifically to *psbA* mRNA, and (ii) light-induced photosynthetic electron transport triggers a plastome-wide increase in translation elongation rate. As a consequence, the synthesis of all plastid-encoded proteins changes rapidly when plants harboring mature chloroplasts are shifted between light and dark, the magnitude of this change is greatest for D1, and the *psbA* mRNA is the only chloroplast mRNA for which large changes in ribosome occupancy accompany the changes in translation rate. The control of *psbA* translation by light-induced D1 damage is of fundamental and practical interest as it maintains functional PSII in the face of inevitable D1 photodamage. It is also a striking example of translational control in both its magnitude and kinetics: the average number of ribosomes per *psbA* transcript decreases more than six-fold within 30 minutes of shifting plants from moderate light intensities to dark and is restored within 15 minutes of shifting plants back to light (Chotewutmontri and Barkan, 2018).

The assembly status of D1 has been implicated in the mechanism underlying the light-regulated recruitment of ribosomes to *psbA* mRNA. This connection began with the findings that the protein HCF244 is required for both the assembly of D1 into the PSII reaction center (Knoppova et al., 2014; Hey and Grimm, 2018; Li et al., 2019; Maeda et al., 2022) and for the recruitment of ribosomes to *psbA* mRNA (Link et al., 2012; Chotewutmontri et al., 2020). Dual functions in D1 assembly and *psbA* translational activation were recently ascribed also to the Arabidopsis protein RBD1 (Garcia-Cerdan et al., 2019; Che et al., 2022; Calderon et al., 2023). Furthermore, the PSII assembly factor HCF136 (Meurer et al., 1998) influences *psbA* translation in a different and particularly revealing way: ribosome occupancy on *psbA* mRNA in maize and Arabidopsis *hcf136* mutants is normal in the light but does not decrease upon a transfer to the dark (Chotewutmontri and Barkan, 2020). HCF136 and its cyanobacterial otholog (Ycf48) act prior to HCF244 during D1’s assembly into the PSII reaction center (Meurer et al., 1998; Plücken et al., 2002; Komenda et al., 2008; Knoppova et al., 2014; Knoppova et al., 2022; Zhao et al., 2023). These results led to a working model in which D1 synthesis is coupled to D1 damage via a translational autoregulatory circuit centered in the HCF244 assembly complex; this model posited that nascent D1 in the HCF244 complex prevents HCF244 from activating *psbA* translation, and that this repression is relieved when D1 is transferred from the HCF244 complex to a D1-less PSII repair intermediate (Chotewutmontri and Barkan, 2020).

These effects on *psbA* translation must be mediated by a direct translational regulator bound to *psbA* mRNA; the protein HCF173 is a strong candidate for such a factor. HCF173 is required specifically for *psbA* translation (Schult et al., 2007; Williams-Carrier et al., 2019) and binds *in vivo* to a site in the *psbA* 5’-untranslated region (UTR) that is adjacent to the footprint of the initiating ribosome (McDermott et al., 2019). Recent results suggest that HCF173 is also necessary to regulate *psbA* translation in response to light (Rojas et al., 2024). Our comprehensive search for proteins that bind *in vivo* to *psbA* mRNA identified several proteins in addition to HCF173, but none of these are required for *psbA* translation (McDermott et al., 2019; Watkins et al., 2020). The pentatricopeptide repeat protein LPE1 had been reported to bind the *psbA* 5’ UTR and to be required for *psbA* translation (Jin et al., 2018), but subsequent reports called those conclusions into question (McDermott et al., 2019; Williams-Carrier et al., 2019; Watkins et al., 2020). Thus, current data strongly suggest that HCF173 is the key regulator at the end of the signal transduction chain that couples D1 photodamage to *psbA* translation initiation.

Experiments described herein began with the goal of defining *cis*-elements in the *psbA* 5’-UTR that mediate the gain and loss of ribosomes on the *psbA* ORF in response to light and D1 damage. For that purpose, we generated reporter constructs in transplastomic tobacco. One construct harbored the *psbA* promoter and 5’-UTR plus the first 22 codons of the *psbA* open reading frame (ORF) fused in-frame to an ORF encoding green fluorescent protein (GFP). We had anticipated that the *psbA* sequences in this construct would be sufficient to regulate GFP synthesis in a manner that recapitulates that of endogenous D1 because translation initiation rates in bacteria and chloroplasts are determined by interactions proximal to the start codon (reviewed in Gualerzi and Pon, 2015; Zoschke and Bock, 2018; Carrier et al., 2020; Willmund et al., 2023). In fact, prior studies had concluded that the *psbA* 5’-UTR is sufficient to program light-regulated translation of reporter genes in tobacco (Staub and Maliga, 1994; Eibl et al., 1999). We show here that the *psbA* 5’ UTR is indeed required for the regulation of *psbA* translation. Contrasting with our expectation, however, the *psbA* UTRs are not sufficient to program light-regulated translation. Instead, we found that the *psbA* ORF plays an essential role, repressing translation initiation in *cis,* with this repression relieved by light and D1 photodamage. We showed further that this repression requires HCF173’s carboxy terminus, revealing that HCF173 itself mediates repression. These results, in conjunction with recent insights into the function of HCF173’s enigmatic “CIA30” domain, lead to a working model positing that co-translational interactions between HCF173 on the *psbA* 5’-UTR and the nascent D1 peptide prevent HCF173 from activating translation, that this repressive interaction is relieved by light-induced D1 damage, and that the PSII assembly factors HCF136 and RBD1 tie D1 synthesis to D1 damage by modulating these interactions.

## RESULTS

### Transplastomic reporter lines for the study of *psbA* translation

RNA coimmunoprecipitation experiments showed that HCF173 associates *in vivo* with a conserved sequence block in the *psbA* 5’ UTR that is adjacent to the footprint of the initiating ribosome (Fig. 1A) (McDermott et al., 2019). We refer to this RNA segment as the HCF173 “footprint” because HCF173 protects it from ribonuclease attack. We initially sought to test how this RNA sequence affects *psbA* translation. To do so, we generated two transplastomic tobacco lines. A control construct, denoted *psbA-gfp,* contained the *psbA* promoter, 5’ UTR and the first 22 *psbA* codons fused in-frame with the *gfp* ORF (Fig. 1B, Supplementary Fig. S1). We included the beginning of the *psbA* ORF because ORF sequences near start codons can influence translation initiation in bacteria and chloroplasts (Kuroda and Maliga, 2001; Jagodnik et al., 2017; Verma et al., 2019; Carrier et al., 2020). The included *psbA* codons end before D1’s first transmembrane segment. The second transplastomic line, *psbA^mut^-gfp* was identical to the first except that it included five point mutations in the HCF173 footprint (Fig. 1A, Supplementary Fig. S1A). Evidence for homoplasmy and phenotypes of the transplastomic lines are shown in Supplementary Figure S2. Our experiments also employed a transplastomic tobacco line created by Koya et al (Koya et al., 2005) that contained a construct similar to *psbA-gfp* except that the *psbA* promoter and UTRs directly flank an ORF encoding anthrax protective antigen (PagA) (Fig. 1C).

**Figure 1.**
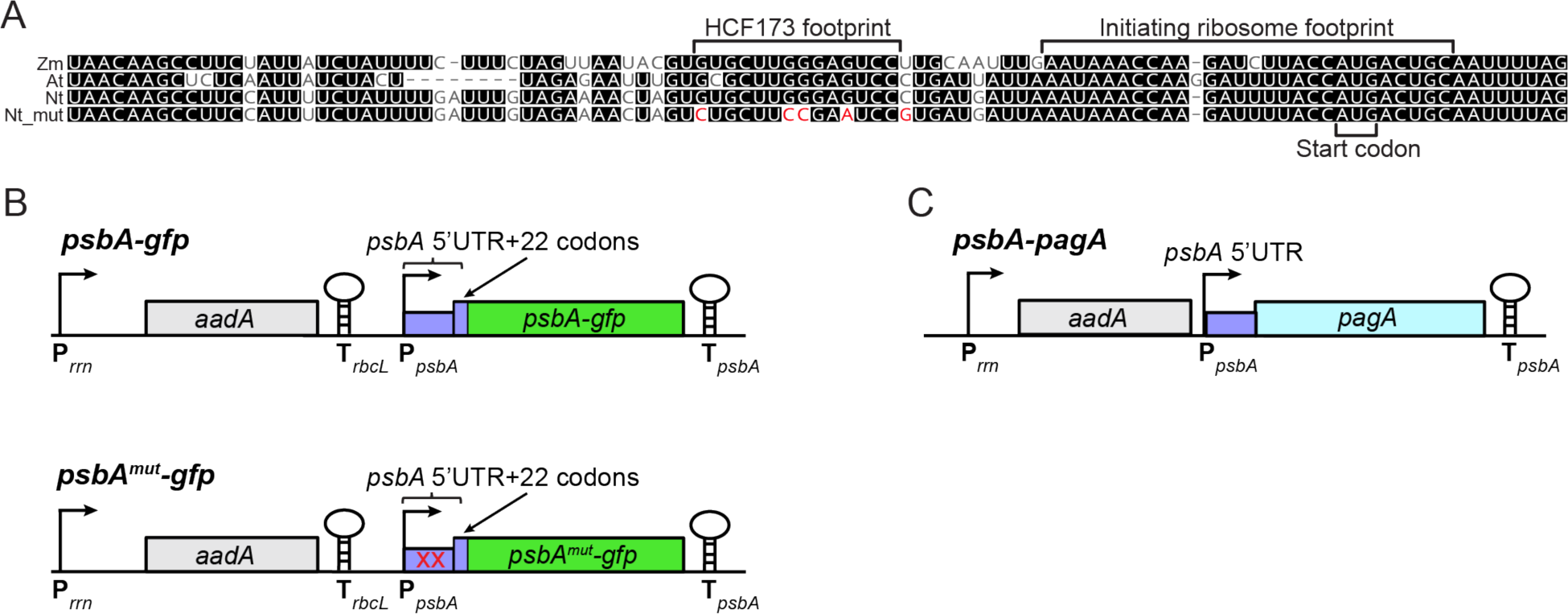
Transplastomic tobacco lines used in this study. (A) Multiple sequence alignment of the *psbA* 5’-UTR in *Zea mays* (Zm), *Arabidopsis thaliana* (At), and *Nicotiana tabacum* (Nt). The *in vivo* footprints of HCF173 (McDermott et al., 2019) and the initiating ribosome (Chotewutmontri and Barkan, 2016) are marked. The mutations incorporated into the *psbA^mut^*-*gfp* construct are highlighted in red (Nt_mut). (B) Chloroplast reporter constructs generated for this study. The selectable marker *aadA* is flanked by the tobacco *rrn* promoter (P*_rrn_*) and the tobacco *rbcL* 3’-UTR (T_rbcL_). In the *psbA-gfp* construct, the tobacco *psbA* promoter, 5’-UTR and first 22 codons of the *psbA* ORF are fused in-frame to the *gfp* ORF, which is followed by the tobacco *psbA* 3’-UTR (T_psbA_). The *psbA^mut^-gfp* construct is identical excepting the five point mutations indicated in panel (A). (C) The *psbA-pagA* construct reported by Koya et al. (Koya et al., 2005). The *pagA* transgene is flanked by the same *psbA* sequences as that in the *psbA-gfp* construct except that the *pagA* ORF was directly adjacent to the *psbA* 5’ UTR.

### The *psbA* UTRs are not sufficient to recapitulate *psbA’*s translational responses to light

We started by using a polysome sedimentation assay to compare effects of light on translation of the *gfp* and *psbA* mRNAs in the *psbA-gfp* reporter line. Leaf tissue from young plants grown in diurnal cycles was harvested after 1 hour of dark adaptation and after 15 minutes of reillumination. Lysates were resolved by ultracentrifugation through sucrose gradients, and RNA extracted from gradient fractions was analyzed by RNA gel blot hybridization (Fig. 2A). The bulk of *psbA* mRNA in the dark-adapted tissue was found in non-polysomal fractions at the top of the gradient and shifted to polysomes after 15 minutes of reillumination, with a substantial fraction remaining non-polysomal even in the light. These results are similar to those reported for *psbA* mRNA in barley, maize, and Arabidopsis (Klein et al., 1988; Chotewutmontri and Barkan, 2018; Gawronski et al., 2021). By contrast, the *gfp* mRNA was largely polysome-associated in both conditions, albeit with a slight reduction in non-polysomal RNA in the reilluminated sample. Similar results were obtained in a replicate experiment using an independent *psbA-gfp* transformant (Supplementary Fig. S3A).

**Figure 2.**
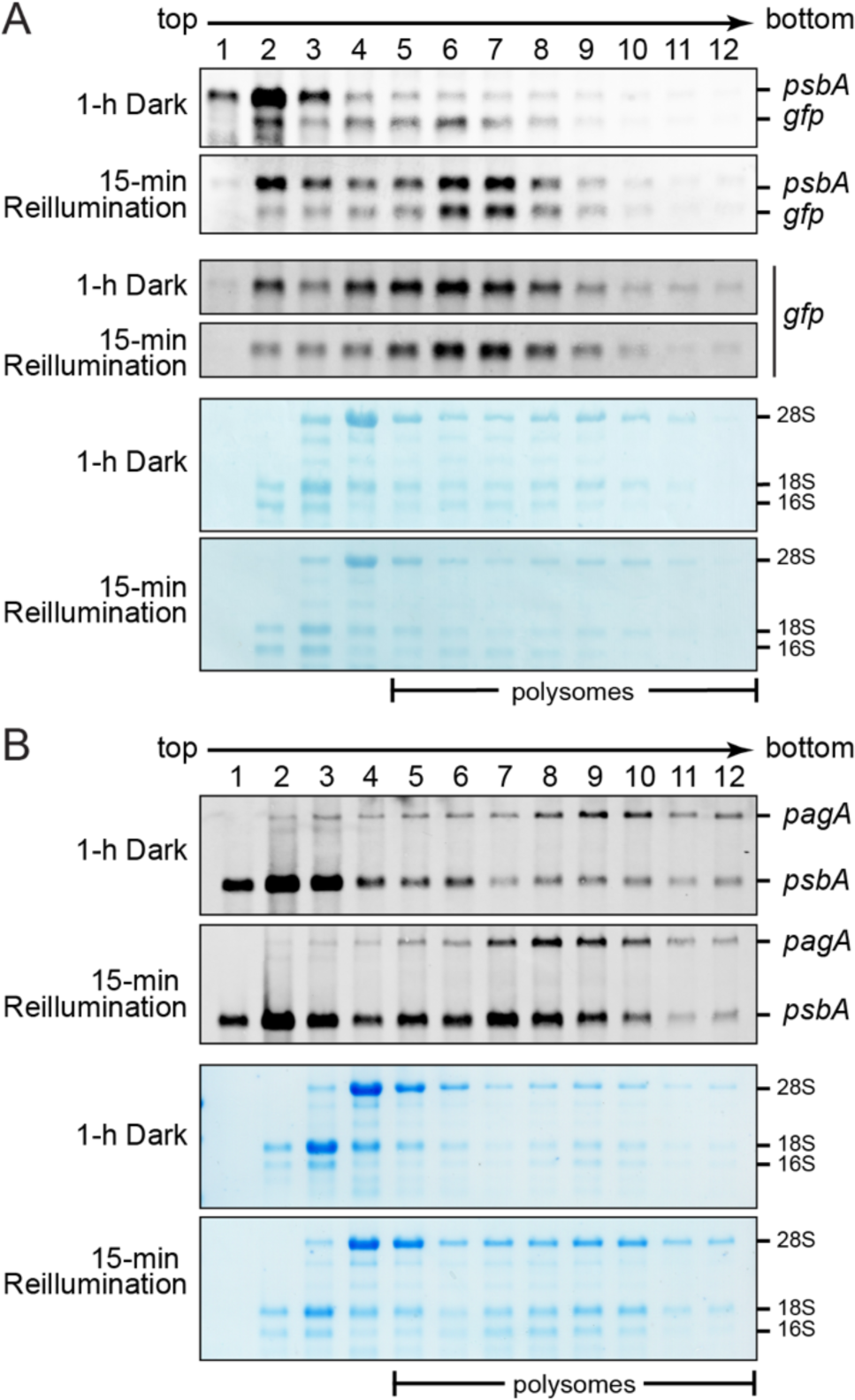
Polysome analysis of reporter mRNAs in dark-adapted and reilluminated leaves. Plants were grown in diurnal cycles for approximately 5 weeks and the youngest fully expanded leaf was harvested after a shift to the dark for 1 hour or after 15 minutes of reillumination. Leaf lysates were sedimented through sucrose gradients, and an equal proportion of the RNA recovered from each fraction was analyzed by RNA gel blot hybridization. Blots were stained with methylene blue prior to probing (below), to show the distribution of rRNAs in the gradient. rRNAs from the cytosol (28S, 18S) and chloroplast (16S) are marked. (A) Polysome analysis of the *psbA-gfp* reporter. The top pair of panels were probed with an oligonucletoide complementary to the first 50 nt of the *psbA* ORF, which is found in both *gfp* and *psbA* mRNAs. To more clearly visualize *gfp* mRNA, the same blots were reprobed with an oligonucleotide specific for *gfp* in the pair of panels below. (B) Polysome analysis of the *pagA* reporter. The blots were probed with an oligonucleotide complementary to the *psbA* 5’ UTR, which is found in both the *pagA* and *psbA* mRNAs.

We also analyzed a previously described transplastomic line in which the *pagA* ORF was flanked by the same *psbA* sequences as those in the *psbA-gfp* construct excepting the *psbA* codons at the start of the ORF (Koya et al., 2005) (see Fig. 1C). The results were similar to those obtained with the *psbA-gfp* reporter: endogenous *psbA* mRNA was largely non-polysomal after one hour in the dark and shifted toward polysome fractions after 15 minutes of reillumination whereas *pagA* mRNA was found in polysomes of similar size in both conditions, albeit with a slight presence in non-polysomal fractions in the dark (Fig. 2B). Similar results were obtained in a replicate experiment (Supplementary Fig. S3B). These results show that the *psbA* UTRs, either on their own or in conjunction with the first 22 *psbA* codons, are not sufficient to recapitulate the effects of light on *psbA* translation.

We showed previously that supplementation of low intensity white light with UVA light specifically induces D1 synthesis and the recruitment of ribosomes to *psbA* mRNA in Arabidopsis (Chotewutmontri and Barkan, 2020). Because UVA induces D1 damage but does not support photosynthetic electron transport (McCree, 1972; Takahashi et al., 2010), this was a key piece of evidence that D1 photodamage is the light-induced signal that triggers the specific increase in *psbA* translation initiation. To determine whether the *psbA* sequences in the *psbA-gfp* and *pagA* constructs are sufficient to program the UVA response, the transplastomic plants were acclimated to low intensity white light for 30 minutes, exposed for 15 minutes to supplemental UVA light, and then pulse-labelled with ^35^S-Met/Cys (Fig. 3A). D1 synthesis was induced by UVA in both the parental and transplastomic lines (Fig. 3B and C), as expected based on results in Arabidopsis (Chotewutmontri and Barkan, 2020). By contrast, synthesis of GFP and PagA proteins did not detectably change.

**Figure 3.**
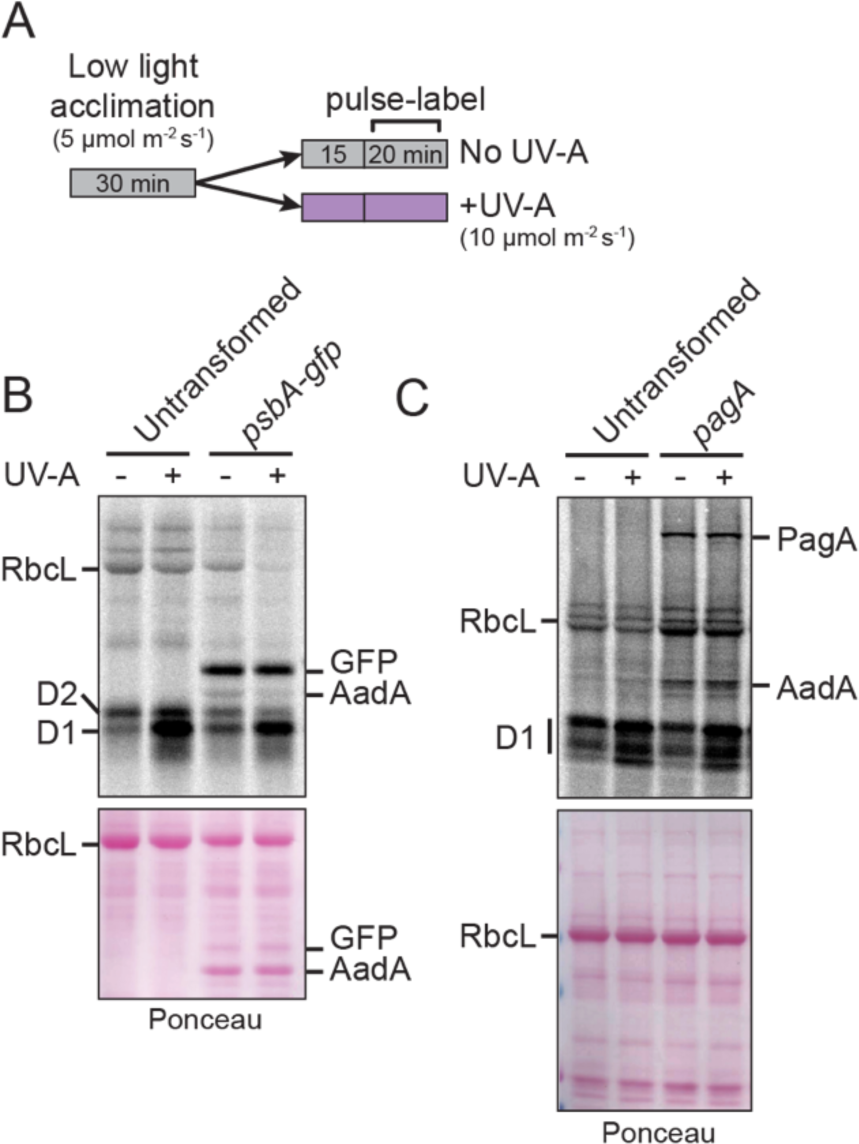
Effect of UVA light on reporter protein synthesis in *psbA* UTR reporter lines. (A) Experimental Design. Plants were grown in soil for five weeks, acclimated to low intensity white light for 30 min and then exposed to supplemental UV-A light (10 µM m^−2^ s^−1^) or maintained in low intensity white light for 15 min. Excised leaf disks were then pulse-labeled with ^35^S-Met/Cys under the same light condition for 20 min. (B) Analysis of the *psbA-gfp* reporter line. Proteins extracted from the pulse-labeled leaves were resolved by SDS-PAGE containing urea, electrophoretically transferred to nitrocellulose and imaged to detect radioactive proteins. The same blot stained with Ponceau S is shown below. (C) Analysis of the *pagA* reporter line. The experiment was performed as described for the *psbA-gfp* line except that the gel did not contain urea.

The polysome and pulse-labeling data demonstrated that the *psbA* UTRs plus the first 22 *psbA* codons are not sufficient to program the translational changes exhibited by *psbA* mRNA in response to light quality and quantity. Instead, sequences downstream from the first 22 codons in the *psbA* ORF are required to prevent ribosome association in the dark, to maintain a ribosome-free fraction of *psbA* mRNA in the light and to regulate D1 synthesis in response to UVA-induced D1 damage. These findings, at first glance, seem inconsistent with conclusions from prior reports (Staub and Maliga, 1993, 1994; Eibl et al., 1999). This apparent inconsistency arises from the term “light regulation” being applied to several entirely distinct phenomena, as discussed below.

### The *psbA* ORF represses translation initiation in *cis* and repression is relieved by light

We next used ribosome profiling to quantify ribosome occupancy on the reporter and *psbA* mRNAs in dark-adapted and illuminated leaves. Ribosome footprints (nuclease-resistant mRNA fragments protected by ribosomes) were purified from leaves of *psbA-gfp* plants harvested at midday, after 1 hour in the dark or after 15 minutes of reillumination. To ensure that minor developmental differences between the sampled leaves did not influence the results, we divided a single leaf along the midrib for the midday versus 1 hour dark comparison, and another leaf for the 1 hour dark versus 15 min reillumination comparison (Fig. 4A). Both comparisons were performed with two biological replicates. The abundance of *psbA* mRNA does not to change during this light shift regime in maize and Arabidopsis seedlings (Chotewutmontri and Barkan, 2018), and RNA gel blot hybridization of RNA purified from lysates used for ribo-seq (Fig. 4B) showed the same to be true for *gfp* and *psbA* mRNAs in *psbA-gfp* tobacco.

**Figure 4.**
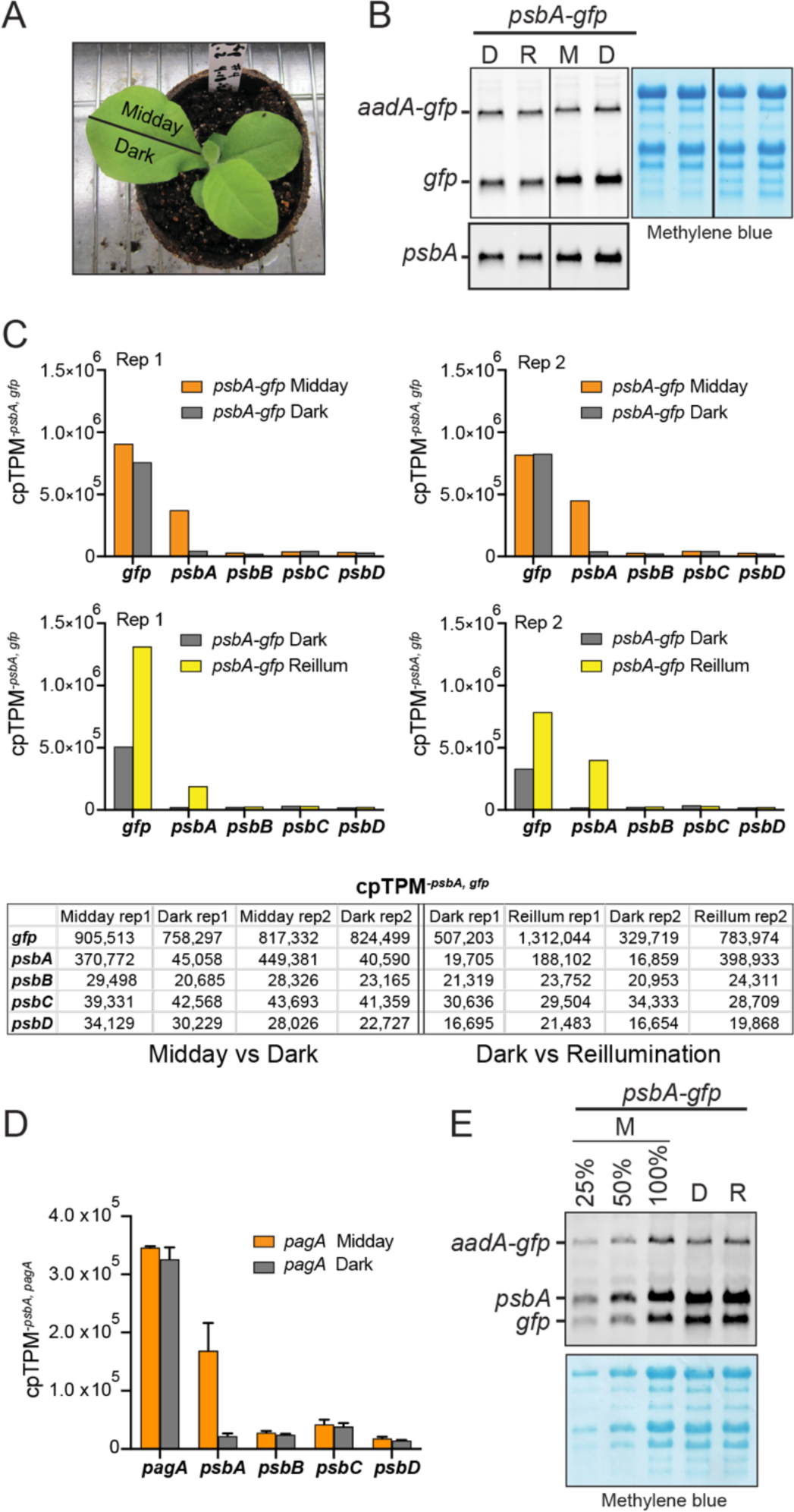
Ribosome profiling analysis of *psbA* UTR reporter lines during light-dark shifts. (A) Tissue used for ribo-seq analysis of *psbA-gfp* plants. A single leaf was divided along the midrib, with tissue harvested at midday or after 1 hour in the dark. The analogous approach was used for comparison of 1 hour of dark adaptation to 15 minutes of reillumination. (B) RNA gel blot analysis of *psbA* and *gfp* mRNAs in lysates of *psbA-gfp* tissue used for ribo-seq. Lanes show RNA from a 1-h dark (D) *versus* 15 min reillumination (R) pair, and from a midday (M) *versus* 1-h dark (D) pair. The blot was hybridized with a probe complementary to the *gfp* ORF (top), and then stripped and hybridized with a probe complementary to a sequence in the *psbA* ORF that is not found in the *gfp* mRNA (bottom). *aadA-gfp* is a dicistronic transcript resulting from read through past the *aadA* gene in the reporter construct (see Fig.1B). The blot was stained with methylene blue (right) to illustrate equal loading of rRNAs. Vertical lines demarcate lanes that were digitally removed from the original image. (C) Ribo-seq analysis of the *psbA-gfp* line. A single divided leaf (see panel A) was used for each of two replicates (Rep) of the Midday versus Dark comparison, and for the Dark versus Reillumination comparison. The abundance of ribosome footprints mapping to each gene was normalized to ORF length and sequencing depth using the cpTPM*^−psbA,gfp^*method. Results for a subset of genes are shown in the graphs and table below. Values for all chloroplast genes are provided in Supplementary Data Set 1. (D) Ribo-seq analysis of the *pagA* line. Leaves at a developmental stage similar to that shown in (A) were harvested from different plants at either midday or after 1-h of dark adaptation. Data were normalized as for the *psbA-gfp* data, except that *psbA* and *pagA* reads were excluded from the read counts used for normalization (cpTPM*^−psbA,pagA^*). The results for all chloroplast genes are provided in Supplementary Data Set 1. The abundance of *pagA* mRNA in the ribo-seq lysates from the dark-adapted and illuminated plants was similar, as quantified by qRT-PCR (Supplementary Fig. S3D). (E) RNA gel blot hybridization to determine relative abundance of *psbA* and *gfp* mRNAs. RNA was extracted from leaves at midday (M), after 1 h in the dark (D), or after 15 min of reillumination (R). The blot was hybridized with a probe that is complementary to the first 50 nucleotides of the *psbA* ORF, which is found in both the *psbA* and *gfp* transcripts. The same blot stained with methylene blue (below) illustrates equal loading of rRNAs.

Ribo-seq read counts mapping to each chloroplast gene in the different datasets were normalized to ORF length and sequencing depth. Sequencing depth normalization used the total number of reads mapping to chloroplast ORFs excluding those mapping to *gfp* and *psbA*. Reads mapping to *psbA* and *gfp* were excluded because they were so abundant (together comprising roughly half of all chloroplast ribosome footprints) that their inclusion artifactually reduces the magnitude of light-induced fluctuations in *gfp* and *psbA* ribosome footprints. Ribosome footprint abundance on chloroplast genes other than *psbA* changes little, if at all, during this light shift regime in maize and Arabidopsis (Chotewutmontri and Barkan, 2018), making that gene set well suited for read depth normalization.

The normalized values for *gfp, psbA* and several other chloroplast genes are shown in Figure 4C. Replicates are displayed in separate graphs so that the data for the adjacent leaf halves that experienced different light conditions can be directly compared. In the midday versus dark comparison (Fig. 4C top), *psbA* experienced a roughly 10-fold decrease in ribosome occupancy following one hour in the dark, whereas ribosome occupancy on endogenous chloroplast ORFs other than *psbA* changed very little (Supplementary Data Set 1), similar to results in maize and Arabidopsis (Chotewutmontri and Barkan, 2018). In stark contrast with *psbA,* ribosome occupancy on the *gfp* ORF was maintained during the one hour in the dark. The dark-*versus*-reilluminated comparisons (Fig. 4C bottom) showed variation in the magnitude of the *psbA* and *gfp* signals in the two replicates, but the trends were nonetheless clear. Ribosome occupancy on *psbA* RNA increased ∼10- and ∼20-fold upon reillumination in replicates 1 and 2, respectively, whereas ribosome occupancies were nearly constant on other endogenous chloroplast genes. The *gfp* reporter displayed an intermediate response, with ribosome occupancy increasing ∼2.5-fold in both replicates. Experiments using leaves from different *psbA-gfp* plants for each light condition gave similar results (Supplementary Fig. S3C) as did a midday-*versus*-dark comparison with the *pagA* reporter line (Fig. 4D). Thus the ribo-seq data confirm and extend the conclusions from the polysome analyses: the *psbA* UTRs plus the first 22 *psbA* codons are not sufficient to program the rapid loss of ribosomes from the *psbA* ORF upon transfer to the dark and are sufficient to program only a small fraction of the increase in *psbA* ribosome occupancy upon reillumination.

To quantitatively compare the average ribosome density on the *gfp* and *psbA* ORFs, we quantified the ratio of *gfp* to *psbA* RNA by probing an RNA gel blot with an oligonucleotide complementary to a *psbA* sequence found in both mRNAs (Fig. 4E); with this method, the relative intensities of the *gfp* and *psbA* mRNA bands reflect the relative abundance of the two mRNAs. Comparison to a dilution series indicated that the combined abundance of the monocistronic *gfp* and dicistronic *aadA-gfp* readthrough transcripts was roughly 75% that of *psbA* mRNA (Fig. 4E). By contrast, ribosome footprints (per kb) from the *gfp* ORF were considerably more abundant than those from the *psbA* ORF in both the light and the dark (Fig. 4C and Supplementary Fig. S3C). Adjusting for the difference in mRNA concentrations, we infer that the average ribosome density on the *gfp* ORF was roughly 3-fold higher than that on the *psbA* ORF at midday, ∼30-fold higher after 1 h in the dark, and ∼4 fold higher after 15 min reillumination. Thus, the *gfp* ORF was much more efficiently translated than the *psbA* ORF under all tested light conditions, but especially in the dark. Given that the *psbA* and *gfp* mRNAs reside in the same chloroplast and share the same UTRs, these results demonstrate that the *psbA* ORF represses translation initiation in *cis,* that this repression is profound in the dark, and that it is relieved (albeit incompletely) by exposure to light.

### Mutation of the HCF173 binding site severely reduces reporter expression and eliminates light-activated translation

We showed previously that a designer RNA binding protein occupying the HCF173 footprint stimulated *psbA* translation in an *hcf173* mutant background but did not restore light-regulated translation (Rojas et al., 2024), suggesting that HCF173 is required for the light response. To further examine HCF173’s role, we analyzed effects of mutating its footprint sequence in the context of the *gfp* reporter. The *psbA^mut^-gfp* construct is identical to *psbA-gfp* except that it includes five point mutations in the HCF173 footprint (Fig. 1). GFP fluorescence was readily detected in plants harboring the *psbA-gfp* construct but not in those with the *psbA^mut^-gfp* construct (Fig. 5A). Immunoblot analysis showed that GFP abundance was roughly 60-fold lower in *psbA^mut^-gfp* leaf tissue than in *psbA-gfp* leaf tissue, whereas HCF173 abundance was not affected (Fig. 5B).

**Figure 5.**
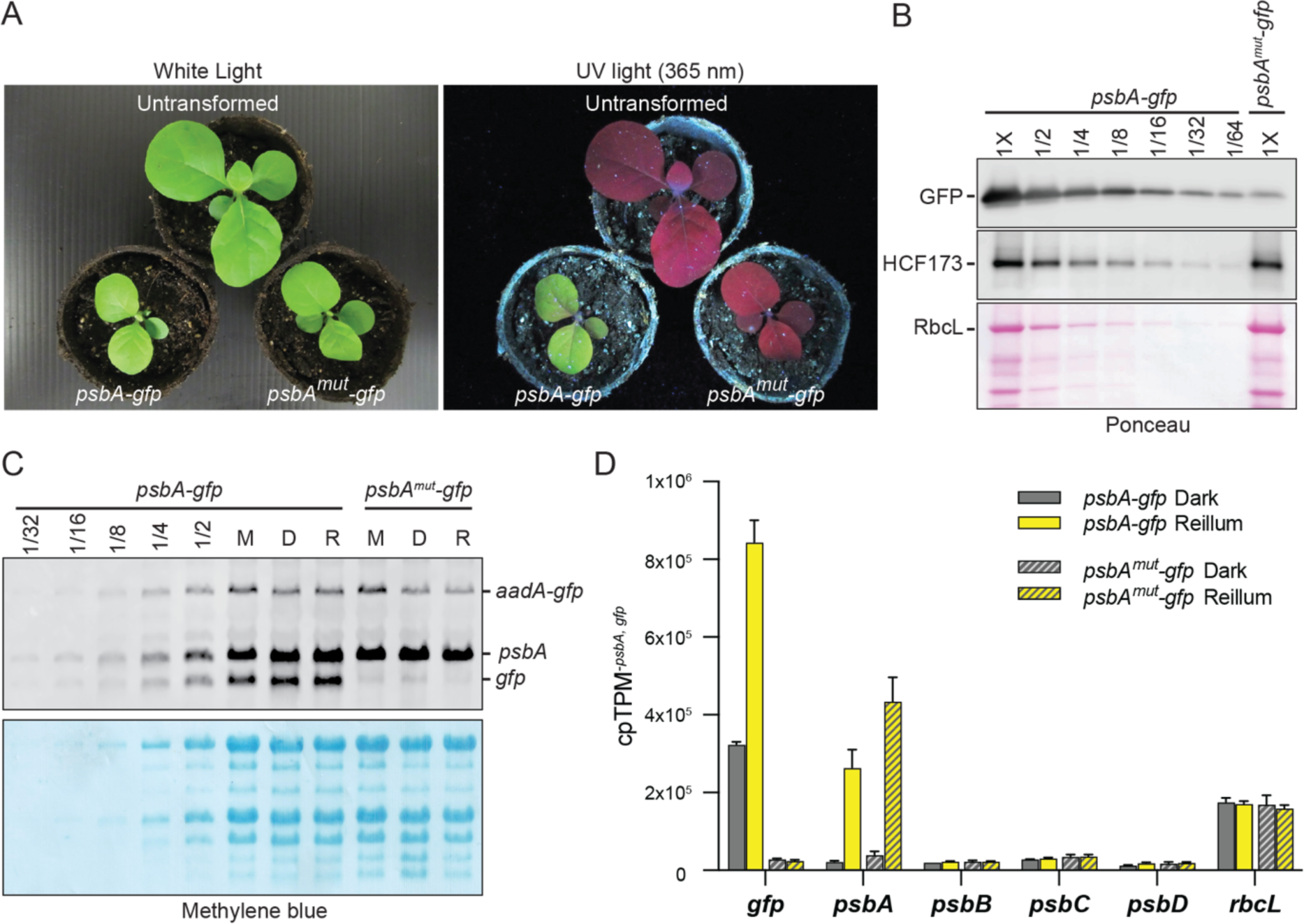
Analysis of *gfp* expression in the *psbA^mut^-gfp* transplastomic line. (A) GFP fluorescence in *psbA-gfp* and *psbA^mut^-gfp* plants. Green and red fluorescence under UV illumination results from GFP and chlorophyll, respectively. (B) Immunoblot quantification of GFP. Total leaf proteins were fractionated by SDS-PAGE, and an immunoblot was probed with antibodies to GFP and HCF173. Total proteins on the same blot were detected by staining with Ponceau S. (C) RNA gel blot hybridization of *gfp* and *psbA* mRNAs using a probe that is complementary to the first 50 nucleotides of the *psbA* ORF, which is found in both the *psbA* and *gfp* transcripts. RNA was extracted from leaves at midday (M), after 1-h in the dark (D), or after 15-min of reillumination (R). Dilutions of the *psbA-gfp* M sample were included to estimate the relative abundance of *gfp* mRNA in the two lines. The same blot stained with methylene blue illustrates equal loading of rRNAs. An excerpt of the same blot is shown in Figure 4E. Results for independent lines are shown in Supplementary Figure S2C. (D) Ribosome profiling analysis of leaf tissue from *psbA-gfp* and *psbA^mut^-gfp* plants harvested after 1-h of dark adaptation or 15-min of reillumination. Values are the mean of two biological replicates +/− SEM. qRT-PCR data confirmed that *gfp* mRNA levels were very similar in the dark-adapted and reilluminated samples from the same line (Supplementary Fig. S3D).

To elucidate the basis of the GFP expression defect in the *psbA^mut^-gfp* line, we analyzed *gfp* mRNA by RNA gel blot hybridization (Fig. 5C). Comparison to a dilution series of RNA from the *psbA-gfp* line showed that monocistronic *gfp* mRNA was roughly 20-fold lower in *psbA^mut^-gfp* leaf tissue whereas the dicistronic *aadA-gfp* read through transcript was similarly abundant in the two lines. The fact that GFP protein in the *psbA^mut^-gfp* line was reduced many-fold more than the sum of its monicistronic and dicistronic mRNAs implies that the mutations also reduced translational efficiency.

It was shown previously that the abundance of *psbA* mRNA in Arabidopsis *hcf173* mutants is reduced roughly 8-fold due to decreased mRNA stability, and that this instability is not a result of the absence of ribosomes on *psbA* mRNA (Schult et al., 2007; Link et al., 2012). Thus, much of the *gfp* mRNA accumulation defect in the *psbA^mut^-gfp* line likely results from the failure of HCF173 to bind and protect a nuclease sensitive site. Nuclease hypersensitivity of the modified RNA sequence might contribute as well. It is curious, however, that the dicistronic *aadA-gfp* read-through transcript was not affected in the *psbA^mut^-gfp* line. Interaction of the upstream RNA with the nuclease-sensitive site might account for this, but the possibility that the HCF173 binding site affects transcription from the *psbA* promoter cannot be excluded.

We next analyzed chloroplast translation in the *psbA^mut^-gfp* line by ribosome profiling, comparing plants that had been dark adapted for 1 hour to those that were reilluminated for 15 minutes (Fig. 5D, Supplementary Data Set 1). The normalized abundance of ribosome footprints on the *gfp* ORF was reduced ∼32-fold in the light and ∼12-fold in the dark by the mutations in the *psbA^mut^-gfp* line. Furthermore, the mutations eliminated the increase in *gfp* ribosome occupancy observed for the *psbA-gfp* reporter when plants were reilluminated. These results show that the RNA sequence occupied by HCF173 *in vivo* is essential for *psbA* expression and suggest that the binding of HCF173 to the *psbA* 5’ UTR is required for the residual light activated translation observed when dark-adapted *psbA-gfp* plants are reilluminated.

### HCF244 and HCF173 activate translation via *cis*-elements in the *psbA* UTRs and do not change in abundance during short-term light shifts

HCF244 is bound to the stromal face of the thylakoid membrane as part of a complex involved in early PSII assembly (Knoppova et al., 2014; Hey and Grimm, 2018; Li et al., 2019; Maeda et al., 2022). Mutation of the *hcf244* gene causes a strong decrease in *psbA* ribosome occupancy in maize and Arabidopsis (Link et al., 2012; Chotewutmontri et al., 2020). However, the mechanism underlying HCF244’s effect on *psbA* translation is unclear. Obvious possibilities could involve interactions with either *psbA* mRNA or with HCF173, but such interactions have not been demonstrated. Given the evidence above that the *psbA* ORF acts in *cis* to regulate *psbA* translation initiation, it seemed plausible that HCF244 exerts its effects on *psbA* translation via the *psbA* ORF rather than via the 5’ UTR.

To address this possibility, we used virus induced gene silencing (VIGS) to knock down *HCF244* and *HCF173* expression in the *psbA-gfp* line. Plants were analyzed one month after VIGS treatment, at which point the abundance of the targeted protein was strongly reduced (Fig. 6A, middle panels). Pulse-labeling assays showed that the synthesis of both D1 and GFP was drastically reduced by VIGS knockdown of either *HCF173* or *HCF244* (Fig. 6A, top). This effect was observed under low intensity white light in both the presence and absence of UVA supplementation. By contrast, the synthesis of other detected proteins was affected little, if at all, by the VIGS treatments. The abundance of *psbA* and *gfp* mRNAs was strongly reduced in the HCF173 knockdown tissue but not in the HCF244 knockdown tissue (Fig. 6B), similar to effects of mutating *HCF173* and *HCF244* on *psbA* mRNA abundance in Arabidopsis (Schult et al., 2007; Link et al., 2012). These results show that both HCF173 and HCF244 activate translation via *cis*-elements that are confined to the *psbA* UTRs plus the first 22 *psbA* codons. These results also show that HCF173’s RNA stabilization effect is mediated by *cis*-elements that are confined to the *psbA* sequences present in the *gfp* mRNA.

**Figure 6.**
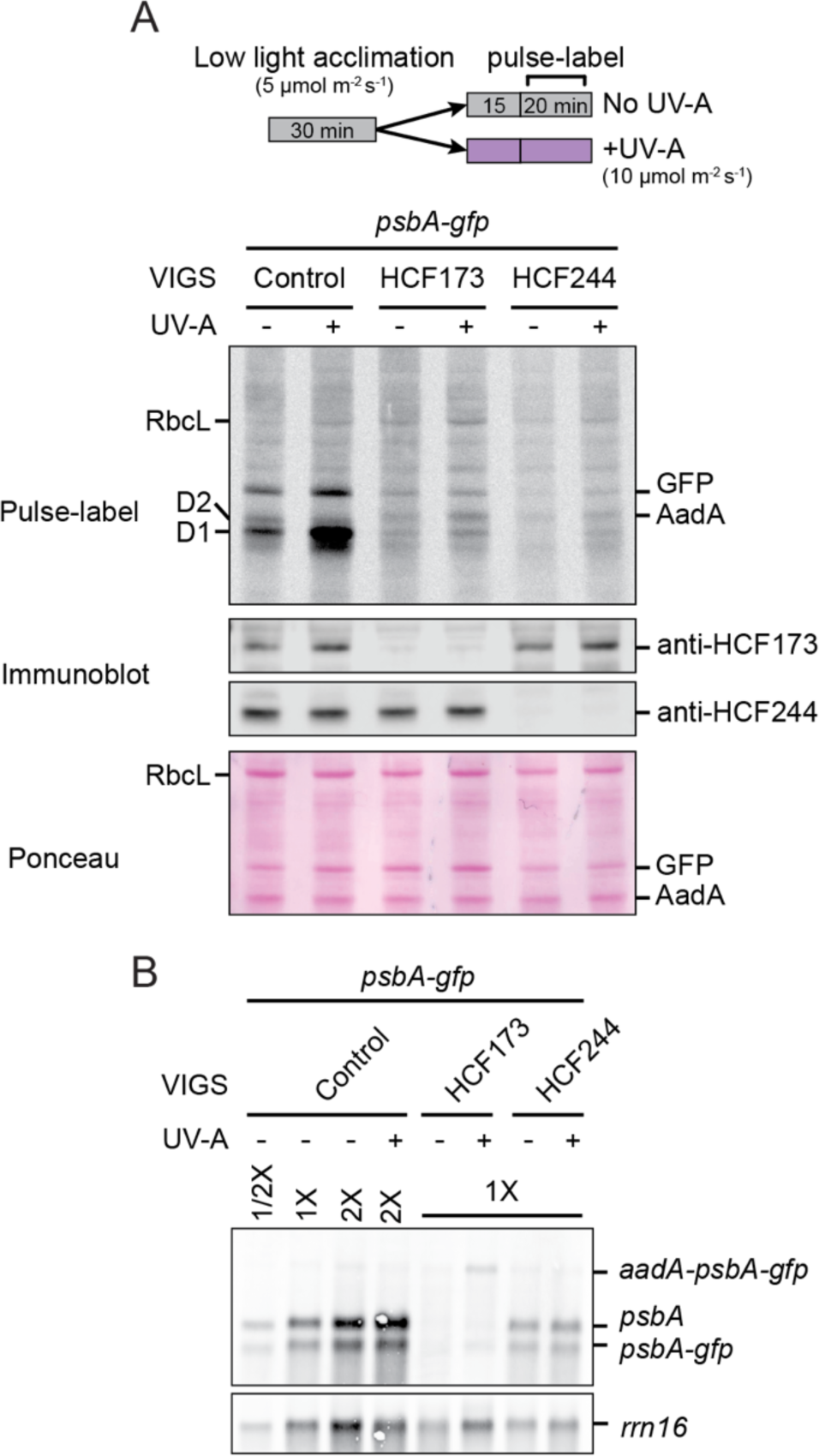
Effects of silencing *HCF173* and *HCF244* on expression of the *psbA-gfp* reporter. (A) Synthesis of D1 and GFP. Control and VIGS-treated *psbA-gfp* plants were acclimated to low intensity white light, and then maintained in that light or exposed to supplemental UV-A for 15 min. Excised leaf disks were then pulse labeled with ^35^S Met/Cys for 20 min in the presence of cycloheximide under the same light condition. Leaf extracts were fractionated in SDS-PAGE gels containing 6M urea, and transferred to nitrocellulose. Radioactive proteins were imaged by phosphorimaging (top panel). Bottom panels show results of probing the same blot with antibodies to either HCF173 or HCF244. An image of the same blot stained with Ponceau S is shown below. AadA is the product of the *aadA* selectable marker in the transplastomic construct. (B) RNA gel blot analysis of *psbA* and *psbA-gfp* mRNA levels in VIGS-treated leaf. The blot was hybridized with an oligonucleotide complementary to a *psbA* sequence present in the *psbA-gfp* transcript (top panel), and then reprobed to detect plastid 16S rRNA (*rrn16*) as a loading control (bottom panel).

We considered the possibility that changes in the abundance of the HCF244 and/or HCF173 contribute to the rapid changes in *psbA* ribosome occupancy when green plants are shifted between light and dark. However, immunoblot analysis showed that their abundance did not change when maize and Arabidopsis seedlings were subject to the light shift regimes used here (Supplementary Fig. S4). This is in accord with prior ribosome profiling data showing that the translational output of the *HCF244* and *HCF173* genes in Arabidopsis did not change when plants were shifted from low intensity to high intensity light (Gawronski et al., 2021), a condition that specifically increases *psbA* translation (Schuster et al., 2019). Therefore, although increased *HCF173* and *HCF244* expression during deetiolation likely contributes to the up-regulation of *psbA* translation during photomorphogenesis (Jin et al., 2018; Kurihara et al., 2020), it does not contribute to the rapid adaptive responses that coordinate D1 synthesis with D1 photodamage.

### The predicted HCF173 structure suggests RNA- and peptide-binding grooves and evokes a potential mechanism for *cis*-repression by the *psbA* ORF

Results above showed that *cis-*elements in the *psbA* 5’ UTR and ORF cooperate to regulate *psbA* translation in response to light. Given that HCF173 acts via the 5’ UTR to activate translation, it seemed plausible that HCF173 integrates information from the *psbA* ORF to program the light response. HCF173 belongs to the short chain dehydrogenase/reductase (SDR) superfamily. Most SDR proteins are oxidoreductases that employ dinucleotide cofactors such as NADP(H) (Persson et al., 2009). However, HCF173 belongs to the “atypical” SDR subfamily (Schult et al., 2007; Link et al., 2012), whose members are thought to be catalytically inactive. Furthermore, HCF173 stands apart from almost all other SDR proteins in that its SDR domain is discontinuous, with a large insertion dividing its SDR domain into two segments (Fig. 7A). The insertion corresponds to a conserved domain denoted CIA30, which was named after its founding member, Complex I Assembly Factor 30 required for the assembly of Complex I in *Neurospora* mitochondria (Küffner et al., 1998). In a structural model of HCF173, HCF173’s two SDR segments assemble into a canonical SDR fold from which the CIA30 domain protrudes (Fig. 7A). The groove separating the CIA30 and SDR domains is lined with basic amino acid residues, suggesting that it may bind RNA (Fig. 7A, bottom).

**Figure 7.**
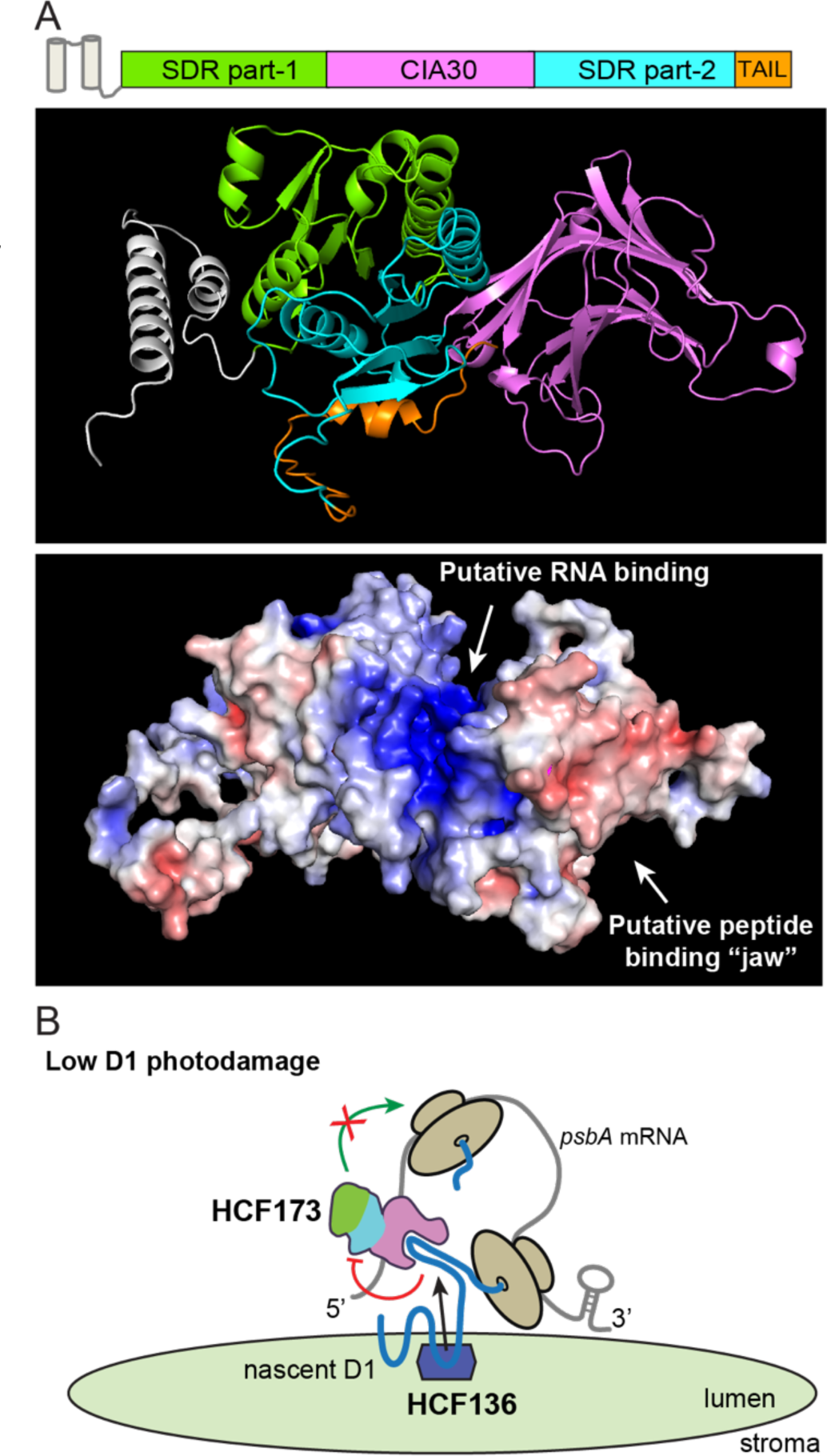
Predicted structure of HCF173 and its implications for mechanisms underlying *cis*-repression of *psbA* translation by the *psbA* ORF. (A) Predicted structure of mature Arabidopsis HCF173 (AtHCF173) lacking its chloroplast targeting peptide. A diagram of HCF173’s domain architecture (top) illustrates its discontinuous SDR domain (green and cyan) interrupted by a CIA 30 domain (magenta), a pair of N-terminal helices (gray) and a C-terminal tail (orange). The structure of AtHCF173 as predicted by AlphaFold is shown below, with domains colored to match the diagram. The electrostatic surface representation rendered with PyMol (bottom) illustrates the basic groove (blue) separating the SDR and CIA30 domains, which we hypothesize binds RNA. The jaw-like feature corresponding to the CIA30/NDUFAF1 region that binds ND3 during Complex I assembly (Schiller et al., 2022) is hypothesized to bind a peptide. The two structural representations are viewed from slightly different angles to best display features of interest. (B) Working model of interactions to account for the repressive effects of HCF136 and the *psbA* ORF on *psbA* translation initiation in conditions of low D1 photodamage. HCF173 (domains colored as above) is bound to the 5’ UTR. The cotranslational binding of nascent D1 to HCF173’s CIA30 jaw is proposed to inhibit HCF173’s ability to activate translation. The inhibitory interaction is proposed to be assisted by HCF136 in the thylakoid lumen, whose cyanobacterial ortholog binds the second lumenal loop of D1 and has been proposed to do so cotranslationally (Yu et al., 2018; Zhao et al., 2023). The model posits that the binding of HCF136 affects the conformation of the opposing stromal segment of D1 in a manner that promotes its inhibitory interaction with HCF173. A more comprehensive model that addresses additional aspects of the phenomenology is presented in Figure 9.

A major advance in understanding the direct biochemical function of CIA30 came with a recent cryo-electron microscopy study of Complex I assembly intermediates (Schiller et al., 2022). In a late assembly intermediate, a jaw-like cleft of the CIA30 ortholog NDUFAF1 binds a matrix-localized loop of the mitochondrially-encoded membrane protein ND3. HCF173’s CIA30 domain is predicted to include the jaw-like feature (Fig. 7A). In analogy to NDUFAF1, it seems reasonable to hypothesize that HCF173’s CIA30 jaw transiently binds a stromal loop of a nascent plastid-encoded membrane protein, potentially D1 itself. An interaction between HCF173 and nascent D1 would provide biochemical connections that could account for several enigmatic features of *psbA* translational control. First, a cotranslational interaction between HCF173’s CIA30 domain and nascent D1 could potentially inhibit HCF173’s ability to activate translation on the mRNA to which it is bound, accounting for the *cis-*repressive effect of the *psbA* ORF (Fig. 7B). A repressive interaction between nascent D1 and HCF173 would also provide the biochemical connections to account for the effects of the PSII assembly factor HCF136 on *psbA* translation. HCF136 is required for the loss of ribosomes on *psbA* mRNA when plants are transferred to the dark (Chotewutmontri and Barkan, 2020), but it resides in the thylakoid lumen so its effect on translation must be indirect. The cyanobacterial HCF136 ortholog Ycf48 binds two lumenal segments of D1, and current data suggest that these interactions are initiated cotranslationally (Yu et al., 2018; Zhao et al., 2023). Thus, HCF136’s interaction with nascent D1 in the lumen could potentially influence the conformation of a stroma-exposed loop of D1 in a manner that facilitates the proposed repressive interaction with HCF173 (Fig. 7B).

### HCF173’s carboxy-terminus is required for repression of *psbA* translation initiation in the dark

A multiple sequence alignment of HCF173 orthologs (Supplementary Fig. S5A) shows strong conservation of the short carboxy-terminal segment mapping downstream of the SDR and CIA30 domains. This segment is predicted to adopt an amphipathic alpha-helix (Supplementary Fig. S5B). This sequence is not conserved in an HCF173 paralog (of unknown function) whose domain organization is otherwise the same as that of HCF173 (Supplementary Fig. S5A). To evaluate its function, we expressed a truncated form of HCF173 lacking this motif (HCF173Trunc) in an Arabidopsis *hcf173* mutant. We appended a carboxy-terminal HA tag to the truncated protein, and expressed the full length HCF173-HA tagged protein as a control. *hcf173* expressing either HCF173-HA or HCF173Trunc-HA grew similarly to Col-0 (Fig. 8A) and accumulated similar levels of D1 (Fig. 8B), implying that both HCF173-HA and HCF173Trunc-HA activate *psbA* expression to a similar degree as wild-type HCF173. However, ribosome profiling analysis showed that both lines were compromised in the repression of *psbA* translation when plants were transferred to the dark. This defect was mild in the HCF173-HA transformants (Fig. 8C), whereas the dark-induced loss of ribosomes from *psbA* mRNA was eliminated in the HCF173Trunc-HA transformants (Fig. 8D). Our previous model proposed HCF244 to be the target of a D1-mediated repressive signal (Chotewutmontri and Barkan, 2020). The constitutively high *psbA* ribosome occupancy conditioned by HCF173Trunc-HA shows that HCF173 transduces a signal that represses translation in the dark, something that has not been proven for HCF244. Therefore, HCF173 is now a stronger candidate for the mediator of a D1-dependent repressive signal. These results add to the evidence that cotranslational repressive interactions between HCF173 and nascent D1 are central to the mechanism that controls *psbA* translation in response to light.

**Figure 8.**
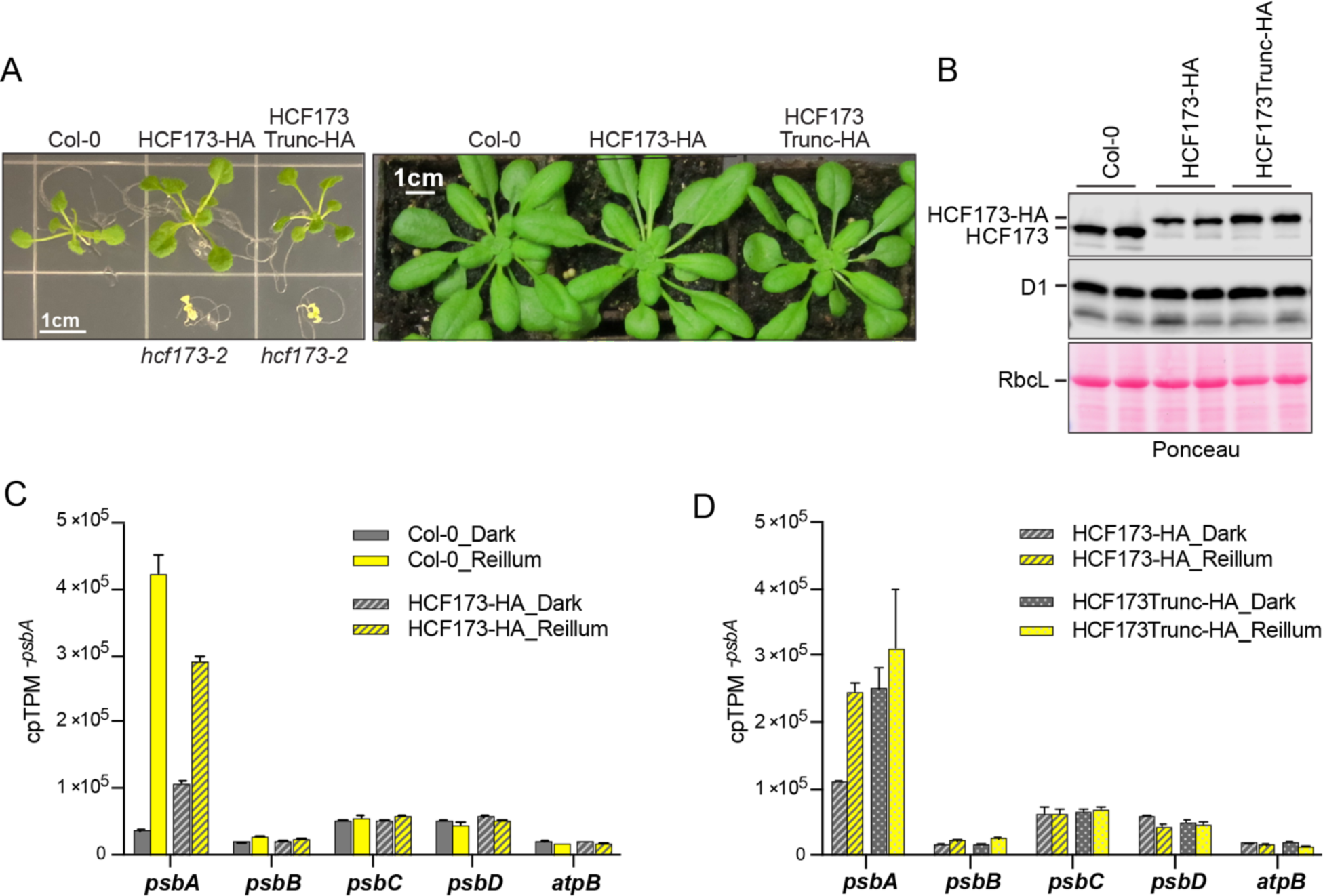
HCF173’s carboxy-terminal tail is required for repression of *psbA* translation initiation in conditions of low D1 damage. A) HCF173-HA and HCF173Trunc-HA complement the visible phenotypes of *hcf173.* Homozygous *hcf173-2* mutants expressing each protein are shown at two developmental stages. Col-0 and *hcf173-2* mutants (which do not survive on soil) are shown for comparison. B) Immunoblot analysis showing the abundance of HCF173, HCF173-HA, HCF173Trunc-HA, and D1 in leaf extracts from the transgenic lines. The blot was probed with an antibody to HCF173 which detects both HA-tagged isoforms. Two seedlings of each genotype were analyzed. The same filter stained with Ponceau S is shown below. C) Ribo-seq data demonstrating compromised dark-repression of *psbA* translation in *hcf173* mutants expressing HCF173-HA. Plants used for ribo-seq were harvested after 1-h of dark adaptation or after 15-min of reillumination. D) Ribo-seq data demonstrating constitutive *psbA* ribosome occupancy in dark-adapted (1-h) or reilluminated (15-min) *hcf173* mutants expressing HCF173Trunc-HA. RNA gel blot data demonstrating similar *psbA* mRNA abundance in these samples is presented in Supplementary Figure S3E.

## DISCUSSION

The activation of D1 synthesis in response to D1 photodamage is an essential and poorly understood aspect of the PSII repair cycle. Results presented here advance understanding of this process in several ways. We show that a repressive *cis*-element in the *psbA* ORF cooperates with an activating *cis*-element in the *psbA* 5’ UTR to regulate translation initiation in response to light and D1 photodamage. *cis-*repression by the *psbA* ORF is essential for the loss of ribosomes from *psbA* mRNA when plants are transferred to the dark, and the carboxy-terminus of the translational activator HCF173 mediates this repression. These findings had not been foreshadowed by prior reports and demand revision of prior models for light-regulated *psbA* translation. In addition, we demonstrate that HCF173 and HCF244 activate translation via *cis-*elements that are confined to the *psbA* UTRs, and we show that the sequence corresponding to HCF173’s footprint in the 5’-UTR is essential for *psbA* expression. The predicted structure of HCF173 together with recent insights into mitochondrial CIA30 suggest that HCF173 binds a plastid-encoded peptide in its CIA30 “jaw”. These observations are discussed below in the context of prior data and incorporated into an updated mechanistic model for light-regulated *psbA* translation.

### *cis-*elements involved in light-regulated *psbA* translation: an update

Our data show that *cis*-elements in the 5’ UTR and ORF cooperate to regulate *psbA* translation in response to light, and that the ORF acts in *cis* to program the loss of ribosomes when plants are shifted to the dark. This role for the ORF may seem at odds with prior reports that the *psbA* 5’ UTR is sufficient to program light-regulated translation of transplastomic reporters in tobacco (Staub and Maliga, 1993, 1994; Eibl et al., 1999). This apparent discrepancy results from the fact that the term “light regulation” has been used to describe several entirely different phenomena. Our focus is on the rapid light-induced changes in *psbA* translation initiation in mature chloroplasts, a phenomenon that is specific to *psbA* and that is triggered by D1 photodamage (Chotewutmontri and Barkan, 2018, 2020). By contrast, the prior reports compared etiolated seedlings grown in the absence of light to seedlings that were then illuminated for several days (Staub and Maliga, 1993, 1994; Eibl et al., 1999). Illumination of etiolated seedlings triggers photomorphogenesis, during which the mRNAs encoding HCF173 and HCF244 are up-regulated (Jin et al., 2018; Kurihara et al., 2020); this likely accounts for the increased translation of *psbA* 5’ UTR reporters when etiolated seedlings are illuminated for extended periods. That phenomenon is distinct from the rapid gain and loss of ribosomes on *psbA* mRNA prompted by shifting green plants between light and dark, during which the abundance of HCF173 and HCF244 does not change (Supplementary Fig. S4).

It was also reported that the *psbA* 5’ UTR is sufficient to program repression of reporter protein synthesis when mature transplastomic plants are shifted to the dark (Staub and Maliga, 1994). However, neither the ribosome association of reporter mRNA nor the expression of reporters with other UTRs was examined. Thus, the drop in reporter protein synthesis likely resulted from the plastome-wide decrease in translation elongation rate that occurs when green plants are shifted to the dark (Chotewutmontri and Barkan, 2018). Our data predict that ribosome occupancy on the reporter changed little, if at all, in that experiment.

### Repressive regulatory interactions underlying light-regulated *psbA* translation

Many chloroplast ORFs require gene-specific nucleus-encoded proteins to be efficiently translated (reviewed in Zoschke and Bock, 2018). The best understood examples activate translation by occupying a 5’ UTR sequence that would otherwise pair with and occlude the ribosome binding site (Klinkert et al., 2006; Prikryl et al., 2011; Hammani et al., 2012; Rahim et al., 2016; Higashi et al., 2021), and this mechanism likely underlies HCF173’s activation of *psbA* translation (McDermott et al., 2019; Gawronski et al., 2021; Rojas et al., 2024). Data presented here and previously (Rojas et al., 2024) show that HCF173 is also essential for the regulation of *psbA* translation in response to light. This regulation could, in principle, involve a signal that activates HCF173 in response to light, a signal that inhibits HCF173 in the absence of light, or both. Our data support the involvement of both positive and negative regulation. A role for negative regulation is indicated by three observations: (i) Mutants lacking the PSII assembly factor HCF136 maintain high *psbA* ribosome occupancy when transferred to the dark (Chotewutmontri and Barkan, 2020); (ii) The *psbA* ORF acts in *cis* to repress translation initiation when plants are transferred to the dark (Figs. 2-4); (iii) Plants expressing truncated HCF173 maintain high *psbA* ribosome occupancy when transferred to the dark (Fig. 8). The working model below invokes a single inhibitory interaction with HCF173 to account for all three of these observations.

Positive regulation of HCF173 in the light is suggested by the observation that *gfp* ribosome occupancy increases several fold when dark-adapted *psbA-gfp* plants are reilluminated (Fig. 4) and that this increase requires the RNA sequence bound by HCF173 (Fig. 5). The combination of these positive and negative regulatory mechanisms could account for the large dynamic range of *psbA* translation initiation in response to changing light intensity.

### Working model for the coupling of *psbA* translation to light-induced D1 damage via cotranslational interactions between nascent D1, HCF173, and HCF136

We proposed previously that *psbA* translation is regulated by D1 photodamage via an autoregulatory circuit centered in the HCF244 PSII assembly complex (Chotewutmontri and Barkan, 2020). The model posited that the presence of D1 in the HCF244 complex prevents HCF244 from communicating with HCF173 to activate *psbA* translation, and that the transfer of D1 to a D1-less PSII repair intermediate relieves that repression. The key observations leading to that model were that HCF173 is a translational activator on the *psbA* 5’UTR, HCF244 is required for both *psbA* translation and PSII assembly/repair, and HCF136, which is required to insert nascent D1 into the HCF244 complex, represses *psbA* translation initiation in the dark. That model predicted that D1 acts in *trans* to repress *psbA* translation in conditions of low D1 photodamage. Repression by D1 is consistent with our new data, but the predicted *trans*-action of D1 is not: the distinct responses of endogenous *psbA* and reporters harboring the *psbA* UTRs shows that the *psbA* ORF acts in *cis* to repress translation initiation when D1 photodamage is low. Furthermore, the previous model does not account for our finding here that HCF173 itself mediates repression of *psbA* translation in the dark, as shown by the constitutively high *psbA* ribosome occupancy in the *HCF173Trunc* mutant. We now propose a revised working model that accounts for these and other recent findings, as well as for the long-recognized coexistence of ribosome-free and actively translated *psbA* mRNA pools.

The model is summarized in Figure 9A,B and illustrated in more detail in Figure 9C,D. In overview, we posit that HCF173 adopts two conformations, both of which bind the *psbA* 5’ UTR but with opposite effects on translation: HCF173^ON^ promotes ribosome binding while HCF173^OFF^ prevents ribosome binding (Fig. 9A). The ratio between the two isoforms varies with the rate of D1 photodamage, thereby changing the partitioning of *psbA* mRNA between translationally active and inactive pools. HCF173^OFF^ prevents translation through steric interference with the binding of HCF173^ON^ to the *psbA* 5’ UTR, and does not itself promote translation, possibly due to steric interference with ribosome binding. When D1 photodamage is low, the HCF173^OFF^ conformation is triggered by HCF136-dependent cotranslational interactions with nascent D1 (Fig. 9B top); once established, repression is maintained after all ribosomes have terminated by the persistent binding of HCF173^OFF^ to the *psbA* 5’ UTR. Repression is maintained until D1 photodamage acts via the PSII assembly factors RBD1 and/or HCF244 to restore the HCF173^ON^ conformation (Fig. 9B, bottom).

**Figure 9.**
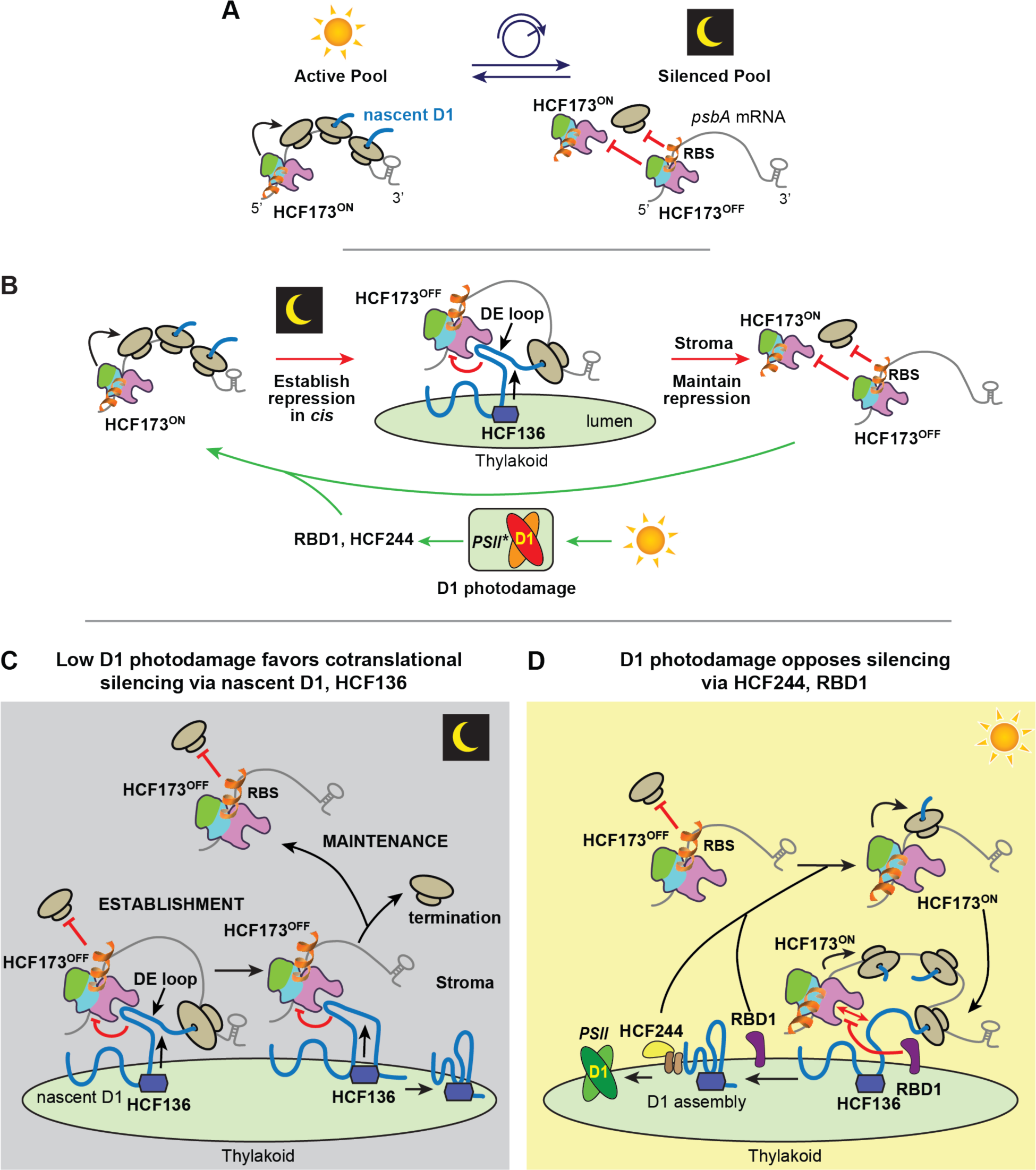
Working model for the regulation of *psbA* translation by light in mature plant chloroplasts. The model is summarized in panels (A) and (B) and is illustrated in more detail in panels (C) and (D). RBS, ribosome binding site. (A) HCF173 adopts two conformations, HCF173^ON^ and HCF173^OFF^, both of which bind the *psbA* 5’UTR. HCF173^OFF^ represses translation by blocking the binding of HCF173^ON^. HCF173’s C-terminal tail is diagrammed as flipping away from the protein core in HCF173^OFF^ such that it sterically blocks the RBS. However, any conformational change that prevents translational activation is plausible. The rate of D1 photodamage shifts the partitioning of *psbA* mRNA between translationally active and inactive pools by modulating the ratio of the two HCF173 isoforms via mechanisms proposed in (B-D). (B) Overview of mechanisms proposed to establish, maintain and reverse the translational silencing of *psbA* mRNA. The HCF173^OFF^ conformation is established cotranslationally via a transient interaction with nascent D1 that is promoted by HCF136 in the thylakoid lumen. Translational silencing is maintained in the stroma by persistent binding of HCF173^OFF^ to the mRNA. Light-induced D1 damage triggers the PSII assembly factors RBD1 and HCF244 to oppose the cotranslational establishment and post-translational maintenance of the HCF173^OFF^ conformation. (C) Establishment and maintenance of repression. A cotranslational interaction between D1’s D-E loop and HCF173’s CIA30 domain is promoted by HCF136, switching HCF173^ON^ to HCF173^OFF^. HCF136 constrains the conformation of D1’s D-E loop in its inhibitory form. The persistent binding of HCF173^OFF^ to the *psbA* 5’ UTR maintains translational repression after ribosomes and nascent D1 are released. This memory of cotranslational silencing events accounts for the coexistence of translationally silenced *psbA* RNA and translationally active *psbA* UTR reporter mRNAs in the stroma. (D) Prevention and reversal of repression. D1 photodamage prevents the establishment of repression by trigging a change in RBD1 activity that opposes the cotranslational inhibitory interaction between D1’s D-E loop and HCF173. D1 photodamage also produces a *trans*-acting signal via HCF244 and/or RBD1 that changes HCF173^OFF^ to HCF173^ON^ on *psbA* mRNAs in the stroma. RBD1 is an integral thylakoid membrane protein that influences the conformation/stability of D1’s D-E loop during PSII assembly/repair and also stimulates *psbA* translation (Garcia-Cerdan et al., 2019; Che et al., 2022; Calderon et al., 2023). HCF244 is tethered to the thylakoid membrane by the integral membrane proteins OHP1 and OHP2, functions in PSII assembly/repair, and is required for *psbA* translation (Link et al., 2012; Li et al., 2019; Hey and Grimm, 2020; Maeda et al., 2022).

The basis for each facet of the model is explained below.

*(i) Repression of psbA translation initiation in conditions of low D1 photodamage is initiated by cotranslational interactions involving nascent D1, HCF173, and HCF136 that promote the HCF173^OFF^ conformation (Fig. 9C).*

Translation initiation in chloroplasts is generally controlled by interactions that modulate the accessibility of the mRNA segment that binds the initiating ribosome (Klinkert et al., 2006; Prikryl et al., 2011; Scharff et al., 2011; Rahim et al., 2016; Scharff et al., 2017; Gawronski et al., 2021; Higashi et al., 2021; Rojas et al., 2024). In that context, two general mechanisms could account for *cis-*repression of translation initiation by the *psbA* ORF: (i) an RNA sequence in the ORF, or a protein that binds such a sequence, establishes a repressive interaction with the ribosome binding site or with a *psbA* translational activator; or (ii) the nascent D1 peptide cotranslationally binds and inhibits a *psbA* translational activator or the ribosome binding site. Observations to date lead us to strongly favor the latter possibility. The fact that the lumen-localized D1 assembly factor HCF136 (Meurer et al., 1998) represses *psbA* translation initiation in the dark (Chotewutmontri and Barkan, 2020) implicates nascent D1 as a mediator of repression. The cyanobacterial HCF136 ortholog binds the lumenal D1 segments flanking its “D” and “E” trans-membrane helices and likely initiates those interactions cotranslationally (Yu et al., 2018; Zhao et al., 2023). This makes D1’s stromal “D-E” loop an attractive candidate for the feature that relays the binding status of HCF136 in the lumen to the stroma (Fig. 9C). Furthermore, the PSII assembly factor RBD1affects the conformation of the D-E loop, and at the same time stimulates *psbA* translation (Garcia-Cerdan et al., 2019; Che et al., 2022; Calderon et al., 2023), supporting the notion that D1’s D-E loop influences *psbA* translation. Translational repression prompted by cotranslational interactions involving nascent D1 provides a plausible explanation for all of these observations.

Finally, the fact that CIA30 binds a matrix-localized loop of a mitochondrially-encoded Complex I subunit during Complex I assembly (Schiller et al., 2022) provides a sound basis for proposing that HCF173’s CIA30 domain binds a stromal loop of D1, and the fact that HCF173 truncation prevents the repression of *psbA* translation initiation in the dark shows that HCF173 itself is the target of a repressive signal. Thus, HCF136-mediated cotranslational repression of HCF173 by nascent D1 - in particular, its D-E loop - provides a parsimonious explanation for current data, providing a biochemical link that connects the *cis* effects of *psbA’s* 5’ UTR and ORF with the *trans* effects of HCF173, HCF136, and RBD1 on *psbA* translation (Fig. 9C).

*(ii) HCF173 is a bistable switch that retains the “OFF” conformation following a transient interaction with nascent D1.*

Ribosome-free *psbA* mRNA is found primarily in the chloroplast stroma (Zoschke and Barkan, 2015; Legen and Schmitz-Linneweber, 2017). The size of this pool increases in conditions of low D1 photodamage, but a substantial ribosome-free pool exists even in moderate growth light (see Fig. 2, for example). A transient interaction between HCF173 and nascent D1 is not sufficient to account for the maintenance of translational repression after the final ribosome on repressed mRNA terminates, at which point the mRNA is released to the stroma. We therefore propose that the HCF173^OFF^ conformation that is prompted by cotranslational interactions with D1 is maintained until an activating signal arises (Fig. 9C). The repressive conformation could be stabilized by, for example, a disulfide bond or the binding of a cofactor.

*(iii) HCF173^OFF^ remains bound to psbA RNA, sterically hindering the binding of HCF173^ON^.*

Our data show that translationally silenced *psbA* mRNA coexists with translationally active *psbA* 5’ UTR reporter mRNAs in conditions of low D1 damage. Given that reporter translation requires HCF173, HCF173^ON^ must be constitutively available in the stroma and yet it does not activate the translationally-silenced *psbA* mRNA pool. To account for this behavior, our model posits that HCF173^OFF^ remains bound to the mRNA on which it was cotranslationally switched off, preventing the binding of HCF173^ON^ by occupying the HCF173 binding site. The fact that *psbA* mRNA abundance does not decline even after 12 hours in the dark in Arabidopsis (Chotewutmontri and Barkan, 2018) supports the proposal that HCF173^OFF^ remains bound to *psbA* mRNA in the dark, because the absence of HCF173 destabilizes *psbA* mRNA. Given the proximity of the HCF173 binding site to the ribosome binding site (Fig. 1A), a small conformational change in HCF173 could also result in steric interference with ribosome binding (Fig. 9A).

*(iv) D1 photodamage prevents cotranslational inhibition of HCF173 and reactivates HCF173^OFF^.*

Illumination prevents ORF-mediated *cis*-repression of *psbA* translation while also reactivating translationally silenced *psbA* mRNA in the stroma. Prior data strongly suggest that both effects are induced by D1 photodamage (Chotewutmontri and Barkan, 2020). We postulate that the D1 assembly factors HCF136 and/or RBD1 couple the rate of D1 photodamage to HCF173’s susceptibility to inhibition by nascent D1 (Fig. 9D). For example, the diversion of HCF136 to PSII repair could prevent HCF136 from promoting the inhibitory interaction between nascent D1 and HCF173, or enhanced activity of RBD1 might influence the longevity of the repressive interaction through its effect on D1’s D-E loop (Calderon et al., 2023).

A distinct mechanism must underlie the reactivation of HCF173^OFF^ in the stroma. Ribosome occupancy increases on *gfp* reporter mRNA when dark-adapted *psbA-gfp* plants are reilluminated; although the magnitude of this increase is considerably less than that on *psbA* mRNA, it demonstrates that a *trans-*activating signal exerts its effect via the *psbA* UTRs. HCF244 is a good candidate for the source of this signal (Fig. 9D) because it is required for translation of the *psbA-gfp* reporter mRNA in the stroma. RBD1 also stimulates *psbA* translation (Che et al., 2022) making it another candidate (Fig. 9D). Both HCF244 and RBD1 are tightly bound to the thylakoid membrane (Link et al., 2012; Garcia-Cerdan et al., 2019; Hey and Grimm, 2020), so it is unlikely that they act directly on silenced stromal *psbA* RNA. RBD1 harbors a redox-active rubredoxin domain that is necessary for PSII repair (Garcia-Cerdan et al., 2019) and mutation of the putative “catalytic tetrad” in HCF244’s SDR domain interfered with D1 synthesis (Hey and Grimm, 2018). Thus, the possibility that a *trans* acting redox signal originating in one or both of these proteins reactivates silenced stromal *psbA* mRNA should be considered.

*(v) D1 synthesis is tuned to D1 photodamage by HCF173-mediated shifts in the partitioning of psbA mRNA between translationally-silenced and translationally active pools.*

In polysome sedimentation experiments, the *psbA* mRNA from plants harvested in the light shows an unusual bimodal distribution, with one peak found in particles that are smaller than ribosomes and the other cosedimenting with polysomes (e.g. Fig. 2 and (Schult et al., 2007). Other chloroplast mRNAs sediment in a single broad peak, with little evidence of non-polysomal RNA (for example Barkan, 1993; Schult et al., 2007). An insufficient quantity of HCF173^ON^ cannot account for the ribosome-free *psbA* mRNA pool because the *psbA-gfp* reporter mRNA, which relies on HCF173 for translation, is almost entirely polysomal in the light. We posit that HCF173^OFF^ is the factor responsible for maintaining the ribosome-free *psbA* mRNA pool (Fig. 9A). The reporter mRNAs escape this silencing, supporting the view that *psbA* mRNAs are drawn into the silenced pool via cotranslational effects of nascent D1. The silenced and active *psbA* mRNA coexist in the stroma, bound to HCF173^OFF^ and HCF173^ON^, respectively, and the partitioning of mRNA molecules between these two pools shifts in response to D1 photodamage (Fig. 9A).

### Relevance to light-regulated *psbA* translation and assembly-coupled translation in Chlamydomonas chloroplasts

A prominent model for light-regulated *psbA* translation in *Chlamydomonas reinhardtii* proposed that photosynthetic electron transport produces a redox signal that enables a translational activator to bind the *psbA* 5’ UTR (Danon and Mayfield, 1994). However, the RNA-protein and protein-protein interactions at the core of that model have not been confirmed with *in vivo* assays. A recent report casts further doubt by showing that disrupting the genes encoding the proposed translational activator and redox-responsive regulator (RB47 and RB60) did not affect PSII activity (Wang et al., 2024). In any case, our data are not compatible with a mechanism of that nature in plants because the activating signal is D1 photodamage, not photosynthetic electron transport (Chotewutmontri and Barkan, 2020), and the *cis*-elements are not confined to the *psbA* 5’ UTR. Chlamydomonas harbors orthologs of all four proteins in our proposed regulatory mechanism (HCF173, HCF244, HCF136, RBD1), and the HCF173, HCF244, and RBD1 orthologs are known to influence PSII biogenesis (Garcia-Cerdan et al., 2019; Kafri et al., 2023; Wang et al., 2024). Therefore, a parsimonious hypothesis is that light regulates *psbA* translation in Chlamydomonas and plant chloroplasts via a similar mechanism.

Our findings have intriguing parallels with assembly-coupled translational controls elucidated in Chlamydomonas chloroplasts (reviewed in Choquet and Wollman, 2023) and yeast mitochondria (Perez-Martinez et al., 2009; Soto et al., 2012; Salvatori et al., 2020), which involve translational repression by unassembled subunits of photosynthetic or respiratory complexes as a means to coordinate subunit synthesis and assembly. In Chlamydomonas chloroplasts, this phenomenon is referred to as “control by epistasis of synthesis” (CES) (Choquet and Wollman, 2023). For example, unassembled D1 represses *psbA* translation in Chlamydomonas, comprising a translational feedback loop to tune D1 synthesis to the need for D1 during PSII assembly (Minai et al., 2006). Defining features of this and other CES examples are that the 5’ UTR of the mRNA subject to translational repression is sufficient to place a reporter gene under CES control, and that the repressing protein acts in *trans* (Choquet and Wollman, 2023); these features also apply to assembly-coupled translation in mitochondria (reviewed in Jung et al., 2024). The mechanism elucidated here likewise connects *psbA* translation and D1 assembly, but the fact that the *psbA* ORF represses translation initiation in *cis* is inconsistent with the CES paradigm. The best understood CES mechanism involves a motif in unassembled cytochrome *f* that triggers the degradation of a translational activator on the 5’ UTR of the mRNA encoding cytochrome *f* (Boulouis et al., 2011). By contrast, the abundance of HCF173 and HCF244 do not change under conditions in which *psbA* translation rate changes, including in mutants with disrupted D1 assembly. Nonetheless, there are strong parallels between CES in Chlamydomonas and light-regulated *psbA* translation in plants, and the differences could be reconciled with small adjustments to the proposed regulatory interactions.

In summary, the findings reported here clarify the mechanism underlying the coupling of D1 photodamage with D1 synthesis for PSII repair. We revealed previously unanticipated roles for the *psbA* open reading frame and for HCF173 in the repression of *psbA* translation in conditions of low D1 photodamage. Our working model suggests plausible intermolecular interactions to account for many features of the phenomenology, including the effects of the PSII assembly factors HCF136 and RBD1 on *psbA* translation and the maintenance of a translationally silent pool of *psbA* mRNA in the stroma. Experiments in progress aim to test each facet of this model and to shed light on aspects that remain particularly enigmatic.

## METHODS

### Tobacco growth and generation of transplastomic tobacco lines

Tobacco (*Nicotiana tabacum* Petit Havana) seeds were plated on agar containing Murashige and Skoog (MS) basal medium (Sigma-Aldrich, MS-5519), 0.6% Micro Agar (Gold Bio) and 3% (w/v) sucrose. When growing transplastomic lines, the medium included 0.5 g/L spectinomycin HCl to select for plants harboring the *aadA* marker. Plants were grown for 2 weeks in a growth chamber at 25°C at a light intensity of 100 µmol m^−2^s^−1^ with 16-h light/8-h dark cycles. Seedlings were then transplanted to soil and grown under the same conditions prior to tissue sampling.

The *pagA* reporter line was generously provided by Henry Daniell (University of Pennsylvania). Our GFP reporter constructs were generated by modification of plasmid pAI3 (Yu et al., 2019), which was generously provided by Pal Maliga and Qiguo Yu (Rutgers University). Sequences of the two *gfp* reporter constructs are provided in Supplementary Figure S1A. Transplastomic plants were generated using the method described in (Maliga et al., 2021). Homoplasmy was assessed by Southern blot hybridization using BamHI to digest 3 µg leaf DNA, and a synthetic DNA oligonucleotide probe complementary to plastid 16S rRNA (Supplementary Table S1), which was 5’-end labelled with infrared dye IRDYE800 (IDT).

### Arabidopsis growth and generation of transgenic Arabidopsis lines

Arabidopsis seeds were sterilized in 30% (v/v) bleach and 0.1% (w/v) SDS for 10 min, washed in 70% (v/v) ethanol, rinsed 3 times in sterile water, and plated on MS medium containing 0.6% (w/v) Micro Agar (Gold Bio) and 3% (w/v) sucrose. Seedlings were grown in a growth chamber in diurnal cycles (10 h light at 80 *µ*mol m^−2^s^−1^, 14 h dark, 22 °C, Philips F25T8/TL841/T8 Fluorescent Bulbs) for 20 days (ribosome profiling) or 16 days (immunoblotting).

The HCF173-HA transgenic line was made by cloning the *HCF173* ORF with a C-terminal 3x Hemagglutinin (HA) tag into pCambia1300 (https://cambia.org/) modified to include the CaMV 35S promoter (enhanced) and the NOS terminator. The ORF sequence was codon-optimized for Arabidopsis. The HCF173Trunc-HA construct was identical except that the sequence encoding the C-terminal 23 amino acids was deleted. Sequences of HCF173-HA and HCF173Trunc-HA are provided in Supplementary Figure S1C. Plants heterozygous for *hcf173-2*, a null allele of *hcf173* (GK-246C02_At1g16720) (Schult et al., 2007), were transformed by the floral dip method (Zhang et al., 2006). Progeny expressing the transgene were identified by resistance to hygromycin and immunoblot analysis using anti-HA antibody. Transformants that were homozygous for *hcf173-2* were identified by PCR using primers flanking the T-DNA insertion, in conjunction with a primer reading outward from the T-DNA left border (Supplementary Table S1). The DNA sequences of each transgene were confirmed in transgenic plants by sequencing PCR products amplified using primers complementary to the CaMV 35S promoter and the sequence encoding the HA tag (Supplementary Table S1). The absence of endogenous HCF173 and presence of HCF173-HA and HCF173Trunc-HA proteins were confirmed by immunoblotting using antibodies to HCF173 and the HA tag, respectively.

### SDS-PAGE and immunoblot analysis

Protein was extracted from leaf tissue, separated by SDS-PAGE and analyzed by immunoblotting as described previously (Barkan, 1998) except for the use of precast Tris-glycine 4-20% polyacrylamide gels (Novex, Invitrogen). The GFP antibody was purchased from Clontech (clone JL-8, 632380) and used at 1:2000 dilution. The AadA antibody was purchased from Agrisera (AS09 580) and used at 1:2000 dilution. The HA antibody was purchased from Agrisera (AS12 2220) and used at 1:5000. The HCF173, HCF244, and D1 antibodies were described previously (McDermott et al., 2019; Chotewutmontri et al., 2020).

### RNA gel blot hybridizations and qRT-PCR

RNA was extracted from leaf tissue with Tri Reagent (Sigma-Aldrich). RNA was analyzed by RNA gel blot hybridization as described previously (Pfalz et al., 2009) except that blots were hybridized with synthetic oligonucleotide probes with IR800 fluorescent tags (IDT). Blots were hybridized overnight in Church Hybridization buffer (7% (w/v) SDS, 0.5 M NaPhosphate pH 7) at 48°C and washed four times for 5 min each in 2XSSC (0.3 M NaCl, 0.03M NaCitrate) and 0.2% (w/v) SDS at 48°C. Results were imaged with a Typhoon Imager (Amersham).

For qRT-PCR, cDNA was prepared by random hexamer priming with Superscript IV (ThermoFisher, 18090010) following the manufacturer’s protocol. qRT-PCR reactions were performed using 5 µl PowerUp SYBR Green Master Mix (Thermo Fisher Scientific, A25742), 1 µl of 2.5 µM forward and reverse primers, 3 µl of water, and 1 µl of diluted cDNA. PCR was performed using a QuantStudio 3 Real Time PCR System (Thermo Fisher Scientific) with initial incubation at 50°C (2 min) followed by 95°C (10 min), and then 40 cycles of 95°C (15 s), 57°C (15 sec) and 60°C (1 min). Each primer set was validated by showing that a single product was amplified in endpoint PCR and by a single peak in melting curve analysis at the end of each qPCR reaction. Each set of assays included four technical replicates and both biological replicates of RNA extracted from ribo-seq lysates. We selected chloroplast *atpB* mRNA as the internal standard because its abundance is more closely matched to that of the chloroplast transgene mRNAs than that from any nuclear gene, and it does not change during the light-shift regimes used here. Primer efficiencies were measured with a standard curve of RNA dilutions, and found to be between 83 and 90% for all primer pairs in the cDNA samples being analyzed. The ΔΔC_T_ relative quantitation method in which fold changes were calculated as 1.85^−ΔΔCT^ to reflect amplification efficiencies of ∼85%.

### Ribosome profiling and polysome analyses

Polysome analyses were performed on the youngest fully expanded leaf from 5 week old tobacco plants, using the method described previously (Barkan, 1998). An equal proportion of the RNA purified from each gradient fraction was analyzed by RNA gel blot hybridization, using probes indicated in each figure.

Ribosome footprints were prepared from the youngest fully expanded leaf from 5 week old tobacco plants or from 3-week old Arabidopsis seedlings using Ambion RNase I (Invitrogen) as described previously (Chotewutmontri et al., 2018). The NEXTflex Small RNA Sequencing Kit v4 (PerkinElmer) was used to generate sequencing libraries. Custom biotinylated antisense oligonucleotides were used for rRNA depletion after first strand cDNA synthesis, as reported previously (Chotewutmontri et al., 2018). Libraries were sequenced using a NovaSeq 6000 or NextSeq 2000 (Illumina) in single-read mode with read lengths of 118 nucleotides.

Ribo-seq data processing was done as described previously (Chotewutmontri and Barkan, 2021) except that trimming did not include removal of four random nucleotides from adapter ends, since the adapters lack these nucleotides. Trimmed reads between 18 and 40 nucleotides were aligned to the tobacco chloroplast genome (GenBank accession Z00044). Reads from transplastomic tobacco were also aligned to the transgene ORFs. Only uniquely-mapping reads that mapped to the correct strand were counted. The reported read counts for each chloroplast gene exclude the first 10 nucleotides of each ORF to avoid counting the pileup of ribosomes near the start codon. In addition, the first 76 nucleotides of the *psbA* ORF was excluded from read counts because that region is found in both endogenous *psbA* and in the *psbA-gfp* transgenes, and the first 55 nucleotides of the atpB ORF was excluded from read counts because that region is found in both *atpB* and in the *aadA* transgene. We report chloroplast-normalized transcript per million (cpTPM) values that were calculated as reads per kilobase, per million reads per kilobase mapped to chloroplast ORFs other than those from the *psbA* and reporter ORFs. The read counts and cpTPM values are provided in Supplementary Data Set 1.

### *In vivo* pulse labeling

Pulse labeling experiments were performed with leaf disks excised from the third or fourth leaf to emerge from tobacco plants that were roughly five weeks old. Plants were first acclimated for 30 min in low intensity (5 μmol m^−2^s^−1^) white light whose spectrum was the same as that of the basal light source used previously (Chotewutmontri and Barkan, 2020). One set of plants was maintained in this light condition, while another set was exposed, in addition, to low intensity (10 μmol m^−2^s^−1^) UVA light. After 15 min, leaf disks (0.6 cm diameter) were excised with a hole punch from the leaf edge near the leaf tip. For each replicate, four leaf disks were placed in a well of a clear 24-well plate containing 135 μL of labeling buffer (10 mM Na-phosphate pH 7.0, 0.1% Tween 20) and wounded by pressing the leaf against the bottom of the well with the tip of a 20-μL pipette tip four times. The plate was placed on top of black paper to avoid reflected light. Labeling was initiated by adding 15 μL of ^35^S-Met/Cys (EasyTag Express ^35^S Protein Labeling Mix, >1000 Ci/mmol, 11 mCi/mL, PerkinElmer). After 20 min, labeling was terminated by rinsing the disks in buffer lacking ^35^S and frozen in liquid N_2_. Lysates were fractionated by SDS-PAGE, electrophoretically transferred to nitrocellulose and imaged with a Storm 825 phosphorimager (GE Healthcare).

### Virus-induced gene silencing

The binary vector system pTRV1 and pTRV2 from tobacco rattle virus (Liu et al., 2002) was used for VIGS. To silence Nt HCF173 or Nt HCF244, we selected 600-bp cDNA fragments from one of the two co-orthologs of each gene (GenBank Accession XM_016652153.1 for Nt HCF244 or XM_016621192.1 for Nt HCF173 fragment). The sequences are provided in Supplementary Figure S1B. These fragments share ≥99% sequence identity between the two co-orthologs. A BLASTN query against the *N. tabacum* genome did not detect other similar sequences. These cDNA fragments were cloned into the BamHI site of pTRV2. *Agrobacterium tumefaciens* GV3101 (pMP90) carrying pTRV1 or pTRV2 derivatives were grown overnight in YEB (5 g/L beef extract, 5 g/L Peptone, 5 g/L sucrose, 1 g/L yeast extract, 0.5 g/L MgSO_4_) supplemented with 50 μg/mL kanamycin, 30 μg/mL gentamicin, and 50 μg/mL rifampicin. Cells were collected by centrifugation at 2,000 x g for 10 min and resuspended in infiltration media (10 mM MES, 20 g/L sucrose, 10 mM MgCl_2_, 200 μM acetosyringone) to O.D._600_ of 3. Cell suspensions carrying pTRV1 and the pTRV2-based plasmids were mixed in a 1:1 ratio (final O.D._600_ of 1.5 for each cell line). *Nicotiana tabacum* Petit Havana was germinated on MS media and transferred to soil in a growth room (16 h at 80 μmol m^−2^ s^−1^ and 8 h dark at 24 °C). Leaf infiltration was performed on the first and second true leaves to emerge, when the leaves were roughly 1 cm in diameter. The mixed cell suspension was infiltrated into the lower leaf using a 1-ml needleless syringe to fill the leaf air space. The infiltrated seedlings were then incubated in diurnal cycles (16 h light, 35 μmol m^−2^ s^−1^ and 8 h dark) at 19 °C for 2 days with a clear plastic cover to maintain moisture, followed by 32 °C for 2 days without cover, and then at 24°C until harvested. Each temperature shift was programmed to change by 1°C/h. Silencing was apparent in roughly one third of the plants as pale green sectors in leaves that emerged subsequent to infiltration. The pale green sectors at ∼1 month post infiltration were used for immunoblotting to confirm the loss of the targeted protein and for pulse-labeling experiments.

**Accession numbers.** The ribo-seq data were submitted to the SRA database under BioProject number PRJNA1177905.

## Acknowledgements

We are grateful to Qiguo Yu (Massachusetts Institute of Technology) for advice about generating transplastomic plants, Pal Maliga (Rutgers University) for providing vector pAI3, Henry Daniell (University of Pennsylvania) for providing the *pagA* transplastomic line, Tessa Burch-Smith (Danforth Center) and Eunsook Park (formerly of UC-Davis) for advice on VIGS, Eunsook Park for VIGS vectors and *Agrobacterium* strains, Reimo Zoschke (Max Planck Institute, Golm) for helpful discussions, Amy Turner (University of Oregon) for help measuring the spectrum of our UV light source, and the University of Oregon Genomics Core Facility for Illumina sequencing. This work benefited from access to the University of Oregon high performance computing cluster, Talapas.

## Author contributions

R.W.-C., P.C., and A.B. designed the research and analyzed the data, R.W.-C., P.C., S.P., M.R. and S.B. performed the research, A.B. wrote the paper, and all authors edited the paper.

## Supplementary data

**Supplementary Figure S1.** Sequences of tobacco transplastomic constructs, Arabidopsis HCF173-HA transgenes, and VIGS constructs

**Supplementary Figure S2.** Characteristics of transplastomic plants

**Supplementary Figure S3.** Data in support of polysome and ribo-seq analyses

**Supplementary Figure S4.** Abundance of HCF244 and HCF173 during short-term light shifts

**Supplementary Figure S5.** Multiple sequence alignment of HCF173 orthologs and paralogs

**Supplementary Table S1.** Primers and nucleic acid probes used in this study

**Supplementary Data Set 1.** Summary of ribo-seq data for chloroplast genes

## Data availability

The data underlying this article are available in the article, in its online supplementary material, and in the SRA database under BioProject PRJNA1177905.

