## Supplemental Data for "Light activates *psbA* translation in plants by relieving inhibition of translation factor HCF173 by the *psbA* ORF in *cis*"

**Supplementary Table S1. DNA oligonucleotides used for PCR and gel blot hybridizations.**

| | |
|---|---|
| | **Probe for DNA gel blot hybridization to evaluate homoplasmy** |
| Nt rrn16 | 5’ ATGGCTGACCGGCGATTACTAGCGATTCCGGCTTCATGCAGGCGAGTTGC |
| | **Probes for RNA gel blot hybridizations** |
| Zm_5’psbAORF | 5’/5IRD800/AAGCGACCCCACAGGCTTGTACTTTCGCGTCTCTCTAAAATTGCAGTCAT |
| Zm_3’psbAORF | 5’/5IRD800/TTATCCATTAAGGTAAGGAACTTCAAGAGCAGCTAGGTCTAGAGGGAAGT |
| Nt-psbA_5’UTR | 5’/5IRD800/TTGGTTTATTTAATCATCAGGGACTCCCAAGCACACTAGTTTTCTACAAA |
| gfp | 5’/5IRD800/GATAATGGTCTGCTAGTTGAACGCTTCCATCTTCAATGTTGTGTCTAATT |
| | **Primers for qRT-PCR** |
| Nt_atpB_F978 | 5' AGTTTATGTACCCGCAGACGAT |
| Nt_atpB_R1123 | 5' GCATGGTTGACGTTGAATCTAA |
| GFP-F115 | 5' GCAACATACGGAAAACTTACCC |
| GFP-R315 | 5'GTCGTCCTTGAAAGAGATGGTC |
| pagA-F_113 | 5'CCCATGGTGGTTACCTCTTCTA |
| pagA-R_303 | 5'TCTTGGTCATCTACCCACATTG |
| | **Primers for PCR to confirm integrity of HCF173-HA and HCF173Trunc-HA transgenes in transgenic plants** |
| CaMV 35S | 5’CACTGACGTAAGGGATGACGC |
| HA tag | 5’CGGGACATCGTAAGGATACGC |
| | **Primers for PCR to identify plants homozygous for *hcf173-2*** |
| At-HCF173-F4 | 5’ACAGACTAGCGCAATTAAGG |
| At-HCF173-R4 | 5’GAATCATTTCACAGAACAAGCTG |
| LB GABI 08409 | 5' ATATTGACCATCATACTCATTGC |

**Supplementary Figure S1. Sequences of tobacco transplastomic constructs, Arabidopsis HCF173-HA transgenes, and VIGS constructs**

**(A) Chloroplast reporter constructs *psbA-gfp* and *psbA$^{mut}$-gfp*.** Features are shaded as follows: (i) arms for homologous recombination (gray), (ii) selectable marker (*rrn* promoter, *atpB* 5' UTR, 14 *atpB* codons, *aadA* ORF, *rbcL* 3' UTR) (yellow), (iii) reporter gene (*psbA* promoter, *psbA* 5' UTR, 22 *psbA* codons, *gfp* ORF, *psbA* 3' UTR) (green). The mutations in the HCF173 binding site in the *psbA$^{mut}$-GFP* are highlighted in red.
**(B) cDNA sequences used for VIGS silencing of *HCF244* and *HCF173* in tobacco.** The Nt*HCF244* sequence came from GenBank Accession XM_016652153.1 and the *NtHCF173* sequence came from XM_016621192.1.
**(C) Sequences encoding HCF173-HA and HCF173Trunc-HA.**

**(A) Chloroplast reporter constructs.**
***psbA-GFP***
AATTCACCGCCGTATGGCTGACCGGCGATTACTAGCGATTCCGGCTTCATGCAGGCGAGTTGCAGCCTGCA
ATCCGAACTGAGGACGGGTTTTTGGGGTTAGCTCACCCTCGCGGGATCGCGACCCTTTGTCCCGGCCATTG
TAGCACGTGTGTCGCCCAGGGCATAAGGGGCATGATGACTTGACGTCATCCTCACCTTCCTCCGGCTTATCA
CCGGCAGTCTGTTCAGGGTTCCAAACTCAACGATGGCAACTAAACACGAGGGTTGCGCTCGTTGCGGGAC
TTAACCCAACACCTTACGGCACGAGCTGACGACAGCCATGCACCACCTGTGTCCGCGTTCCCGAAGGCAC
CCCTCTCTTTCAAGAGGATTCGCGGCATGTCAAGCCCTGGTAAGGTTCTTCGCTTTGCATCGAATTAAACCA
CATGCTCCACCGCTTGTGCGGGCCCCCGTCAATTCCTTTGAGTTTCATTCTTGCGAACGTACTCCCCAGGC
GGGATACTTAACGCGTTAGCTACAGCACTGCACGGGTCGATACGCACAGCGCCTAGTATCCATCGTTTACGG
CTAGGACTACTGGGGTATCTAATCCCATTCGCTCCCCTAGCTTTCGTCTCTCAGTGTCAGTGTCGGCCCAGC
AGAGTGCTTTCGCCGTTGGTGTTCTTTCCGATCTCTACGCATTTCACCGCTCCACCGGAAATTCCCTCTGCC
CCTACCGTACTCCAGCTTGGTAGTTTCCACCGCCTGTCCAGGGTTGAGCCCTGGGATTTGACGGCGGACTT
AAAAAGCCACCTACAGACGCTTTACGCCCAATCATTCCGGATAACGCTTGCATCCTCTGTATTACCGCGGCT
GCTGGCACAGAGTTAGCCGATGCTTATTCCCCAGATACCGTCATTGCTTCTTCTCCGGGAAAAGAAGTTCAC
GACCCGTGGGCCTTCTACCTCCACGCGGCATTGCTCCGTCAGGCTTTCGCCCATTGCGGAAAATTCCCCAC
TGCTGCCTCCCGTAGGAGTCTGGGCCGTGTCTCAGTCCCAGTGTGGCTGATCATCCTCTCGGACCAGCTAC
TGATCATCGCCTTGGTAAGCTATTGCCTCACCAACTAGCTAATCAGACGCGAGCCCCTCCTCGGGCGGATTC
CTCCTTTTGCTCCTCAGCCTACGGGGTATTAGCAGCCGTTTCCAGCTGTTGTTCCCCTCCCAAGGGCAGGT
TCTTACGCGTTACTCACCCGTCCGCCACTGGAAACACCACTTCCCGTCCGACTTGCATGTGTTAAGCATGCC
GCCAGCGTTCATCCTGAGCCAGGATCGAACTCTCCATGAGATTCATAGTTGCATTACTTATAGCTTCCTTGTT
CGTAGACAAAGCGGATTCGGAATTGTCTTTCATTCCAAGGCATAACTTGTATCCATGCGCTTCATATTCGCCC
GGAGTTCGCTCCCAGAAATATAGCCATCCCTGCCCCCTCACGTCAATCCCACGAGCCTCTTATCCATTCTCAT
TGAACGACGGCGGGGGAGCTTTCGAGGCCTCGAAATCCAACTAGAAAAACTCACATTGGGCTTAGGGATAA
TCAGGCTCGAACTGATGACTTCCACCACGTCAAGGTGACACTCTACCGCTGAGTTATATCCCTTCCCCGCCC
CATCGAGAAATAGAACTGACTAATCCTAAGTCAAAGGGTCGAGAAACTCAACGCCACTATTCTTGAACAACTT
GGAGCCGGGCCTTCTTTTCGCACTATTACGGATATGAAAATAATGGTCAAAATCGGATTCAATTGTCAAAGTA
CTTTTGGAATTCGAGCTCGCTCCCCCGCCGTCGTTCAATGAGAATGGATAAGAGGCTCGTGGGATTGACGT
GAGGGGGCAGGGATGGCTATATTTCTGGGAGAATTAACCGATCGACGTGCAAGCGGACATTTATTTTAAATTC
GATAATTTTTGCAAAAACATTTCGACATATTTATTTATTTTATTATTATGAGAATCAATCCTACTACTTCTGGTTCT
GGGGTTTCCACGGCTACTAGCGAAGCGGTGATCGCCGAAGTATCGACTCAACTATCAGAGGTAGTTGGCGT
CATCGAGCGCCATCTCGAACCGACGTTGCTGGCCGTACATTTGTACGGCTCCGCAGTGGATGGCGGCCTG
AAGCCACACAGTGATATTGATTTGCTGGTTACGGTGACCGTAAGGCTTGATGAAACAACGCGGCGAGCTTTG
ATCAACGACCTTTTGGAAACTTCGGCTTCCCCTGGAGAGAGCGAGATTCTCCGCGCTGTAGAAGTCACCAT
TGTTGTGCACGACGACATCATTCCGTGGCGTTATCCAGCTAAGCGCGAACTGCAATTTGGAGAATGGCAGC
GCAATGACATTCTTGCAGGTATCTTCGAGCCAGCCACGATCGACATTGATCTGGCTATCTTGCTGACAAAAG
CAAGAGAACATAGCGTTGCCTTGGTAGGTCCAGCGGCGGAGGAACTCTTTGATCCGGTTCCTGAACAGGAT
CTATTTGAGGCGCTAAATGAAACCTTAACGCTATGGAACTCGCCGCCCGACTGGGCTGGCGATGAGCGAAA
TGTAGTGCTTACGTTGTCCCGCATTTGGTACAGCGCAGTAACCGGCAAAATCGCGCCGAAGGATGTCGCTG
CCGACTGGGCAATGGAGCGCCTGCCGGCCCAGTATCAGCCCGTCATACTTGAAGCTAGACAGGCTTATCTT
GGACAAGAAGAAGATCGCTTGGCCTCGCGCGCAGATCAGTTGGAAGAATTTGTCCACTACGTGAAAGGCGA
GATCACCAAGGTAGTGGGCAAAGAACAAAAACTCATTTCTGAAGAAGACTTGTAACTGCAGAGACATTAGCA
GATAAATTAGCAGGAAATAAAGAAGGATAAGGAGAAAGAACTCAAGTAATTATCCTTCGTTCTCTTAATTGAAT
TGCAATTAAACTCGGCCCAATCTTTTACTAAAAGGATTGAGCCGAATACAACAAAGATTCTATTGCATATATTTT

GACTAAGTATATACTTACCTAGATATACAAGATTTGAAATACAAAATCTAACATACACCTTGGTTGACACGAGTA
TATAAGTCATGTTATACTGTTGAATAACAAGCCTTCCATTTTCTATTTTGATTTGTAGAAAACTAGTGTGCTTGG
GAGTCCCTGATGATTAAATAAACCAAGATTTTACCATGACTGCAATTTTAGAGAGACGCGAAAGCGAAAGCCT
ATGGGGTCGCTTCTGTAACTGGATAACTGCTAGCAGTAAAGGAGAAGAACTTTTCACTGGAGTTGTCCCAAT
TCTTGTTGAATTAGATGGTGATGTTAATGGGCACAAATTTTCTGTCAGTGGAGAGGGTGAAGGTGATGCAAC
ATACGGAAAACTTACCCTTAAATTTATTTGCACTACTGGAAAACTACCTGTTCCTTGGCCAACACTTGTCACTA
CTTTCTCTTATGGTGTTCAATGCTTTTCAAGATACCCAGATCATATGAAGCGGCACGACTTCTTCAAGAGCGC
CATGCCTGAGGGATACGTGCAGGAGAGGACCATCTCTTTCAAGGACGACGGGAACTACAAGACACGTGCT
GAAGTCAAGTTTGAGGGAGACACCCTCGTCAACAGGATCGAGCTTAAGGGAATCGATTTCAAGGAGGACGG
AAACATCCTCGGCCACAAGTTGGAATACAACTACAACTCCCACAACGTATACATCACGGCAGACAAACAAAA
GAATGGAATCAAAGCTAACTTCAAAATTAGACACAACATTGAAGATGGAAGCGTTCAACTAGCAGACCATTAT
CAACAAAATACTCCAATTGGCGATGGCCCTGTCCTTTTACCAGACAACCATTACCTGTCCACACAATCTGCCC
TTTCGAAAGATCCCAACGAAAAGAGAGACCACATGGTCCTTCTTGAGTTTGTAACAGCTGCTGGGATTACAC
ATGGCATGGATGAACTATACAAATAAGCTCTAGCTAGAGCGATCCTGGCCTAGTCTATAGGAGGTTTTGAAAA
GAAAGGAGCAATAATCATTTTCTTGTTCTATCAAGAGGGTGCTATTGCTCCTTTCTTTTTTTCTTTTTATTTATTT
ACTAGTATTTTACTTACATAGACTTTTTTGTTTACATTATAGAAAAAGAAGGAGAGGTTATTTTCTTGCATTTATT
CATGGGGGATCAAAGCTTGATACTTTAACTGCCCCTATCGGAAATAGGATTGACTACCGATTCCGAAGGAACT
GGAGTTACATCTCTTTTCCATTCAAGAGTTCTTATGCGTTTCCACGCCCCTTTGAGACCCCGAAAAATGGACA
AATTCCTTTTCTTAGGAACACATACAAGATTCGTCACTACAAAAAGGATAATGGTAACCCTACCATTAACTACTT
CATTTATGAATTTCATAGTAATAGAAATACATGTCCTACCGAGACAGAATTTGGAACTTGCTATCCTCTTGCCTA
GCAGGCAAAGATTTACCTCCGTGGAAAGGATGATTCATTCGGATCGACATGAGAGTCCAACTACATTGCCAG
AATCCATGTTGTATATTTGAAAGAGGTTGACCTCCTTGCTTCTCTCATGGTACACTCCTCTTCCCGCCGAGCC
CCTTTTCTCCTCGGTCCACAGAGACAAAATGTAGGACTGGTGCCAACAATTCATCAGACTCACTAAGTCGGG
ATCACTAACTAATACTAATCTAATATAATAGTCTAATATATCTAATATAATAGAAAATACTAATATAATAGAAAAGAAC
TGTCTTTTCTGTATACTTTCCCCGGTTCCGTTGCTACCGCGGGCTTTACGCAATCGATCGGATTAGATAGATAT
CCCTTCAACATAGGTCATCGAAAGGATCTCGGAGACCCACCAAAGTACGAAAGCCAGGATCTTTCAGAAAAC
GGATTCCTATTCAAAGAGTGCATAACCGCATGGATAAGCTCACACTAACCCGTCAATTTGGGATCCAAATTCG
AGATTTTCCTTGGGAGGTATCGGGAAGGATTTGGAATGGAATAATATCGATTCATACAGAAGAAAAGGTTCTC
TATTGATTCAAACACTGTACCTAACCTATGGGATAGGGATCGAGGAAGGGGAAAAACCGAAGATTTCACATGG
TACTTTTATCAATCTGATTTATTTCGTACCTTTCGTTCAATGAGAAAATGGGTCAAATTCTACAGGATCAAACCT
ATGGGACTTAAGGAATGATATAAAAAAAAGAGAGGGAAAATATTCATATTAAATAAATATGAAGTAGAAGAACCC
AGATTCCAAATGAACAAATTCAAACTTGAAAAGGATCTTCCTTATTCTTGAAGAATGAGGGGCAAAGGGATTG
ATCAAGAAAGATC

***psbA$^{mut}$-GFP***
AATTCACCGCCGTATGGCTGACCGGCGATTACTAGCGATTCCGGCTTCATGCAGGCGAGTTGCAGCCTGCA
ATCCGAACTGAGGACGGGTTTTTGGGGTTAGCTCACCCTCGCGGGATCGCGACCCTTTGTCCCGGCCATTG
TAGCACGTGTGTCGCCCAGGGCATAAGGGGCATGATGACTTGACGTCATCCTCACCTTCCTCCGGCTTATCA
CCGGCAGTCTGTTCAGGGTTCCAAACTCAACGATGGCAACTAAACACGAGGGTTGCGCTCGTTGCGGGAC
TTAACCCAACACCTTACGGCACGAGCTGACGACAGCCATGCACCACCTGTGTCCGCGTTCCCGAAGGCAC
CCCTCTCTTTCAAGAGGATTCGCGGCATGTCAAGCCCTGGTAAGGTTCTTCGCTTTGCATCGAATTAAACCA
CATGCTCCACCGCTTGTGCGGGCCCCCGTCAATTCCTTTGAGTTTCATTCTTGCGAACGTACTCCCCAGGC
GGGATACTTAACGCGTTAGCTACAGCACTGCACGGGTCGATACGCACAGCGCCTAGTATCCATCGTTTACGG
CTAGGACTACTGGGGTATCTAATCCCATTCGCTCCCCTAGCTTTCGTCTCTCAGTGTCAGTGTCGGCCCAGC
AGAGTGCTTTCGCCGTTGGTGTTCTTTCCGATCTCTACGCATTTCACCGCTCCACCGGAAATTCCCTCTGCC
CCTACCGTACTCCAGCTTGGTAGTTTCCACCGCCTGTCCAGGGTTGAGCCCTGGGATTTGACGGCGGACTT
AAAAAGCCACCTACAGACGCTTTACGCCCAATCATTCCGGATAACGCTTGCATCCTCTGTATTACCGCGGCT
GCTGGCACAGAGTTAGCCGATGCTTATTCCCCAGATACCGTCATTGCTTCTTCTCCGGGAAAAGAAGTTCAC
GACCCGTGGGCCTTCTACCTCCACGCGGCATTGCTCCGTCAGGCTTTCGCCCATTGCGGAAAATTCCCCAC
TGCTGCCTCCCGTAGGAGTCTGGGCCGTGTCTCAGTCCCAGTGTGGCTGATCATCCTCTCGGACCAGCTAC
TGATCATCGCCTTGGTAAGCTATTGCCTCACCAACTAGCTAATCAGACGCGAGCCCCTCCTCGGGCGGATTC
CTCCTTTTGCTCCTCAGCCTACGGGGTATTAGCAGCCGTTTCCAGCTGTTGTTCCCCTCCCAAGGGCAGGT
TCTTACGCGTTACTCACCCGTCCGCCACTGGAAACACCACTTCCCGTCCGACTTGCATGTGTTAAGCATGCC
GCCAGCGTTCATCCTGAGCCAGGATCGAACTCTCCATGAGATTCATAGTTGCATTACTTATAGCTTCCTTGTT
CGTAGACAAAGCGGATTCGGAATTGTCTTTCATTCCAAGGCATAACTTGTATCCATGCGCTTCATATTCGCCC
GGAGTTCGCTCCCAGAAATATAGCCATCCCTGCCCCCTCACGTCAATCCCACGAGCCTCTTATCCATTCTCAT
TGAACGACGGCGGGGGAGCTTTCGAGGCCTCGAAATCCAACTAGAAAAACTCACATTGGGCTTAGGGATAA
TCAGGCTCGAACTGATGACTTCCACCACGTCAAGGTGACACTCTACCGCTGAGTTATATCCCTTCCCCGCCC

CATCGAGAAATAGAACTGACTAATCCTAAGTCAAAGGGTCGAGAAACTCAACGCCACTATTCTTGAACAACTT
GGAGCCGGGCCTTCTTTTCGCACTATTACGGATATGAAAATAATGGTCAAAATCGGATTCAATTGTCAAAGTA
CTTTTGGAATTCGAGCTCGCTCCCCCGCCGTCGTTCAATGAGAATGGATAAGAGGCTCGTGGGATTGACGT
GAGGGGGCAGGGATGGCTATATTTCTGGGAGAATTAACCGATCGACGTGCAAGCGGACATTTATTTTAAATTC
GATAATTTTTGCAAAAACATTTCGACATATTTATTTATTTTATTATTATGAGAATCAATCCTACTACTTCTGGTTCT
GGGGTTTCCACGGCTACTAGCGAAGCGGTGATCGCCGAAGTATCGACTCAACTATCAGAGGTAGTTGGCGT
CATCGAGCGCCATCTCGAACCGACGTTGCTGGCCGTACATTTGTACGGCTCCGCAGTGGATGGCGGCCTG
AAGCCACACAGTGATATTGATTTGCTGGTTACGGTGACCGTAAGGCTTGATGAAACAACGCGGCGAGCTTTG
ATCAACGACCTTTTGGAAACTTCGGCTTCCCCTGGAGAGAGCGAGATTCTCCGCGCTGTAGAAGTCACCAT
TGTTGTGCACGACGACATCATTCCGTGGCGTTATCCAGCTAAGCGCGAACTGCAATTTGGAGAATGGCAGC
GCAATGACATTCTTGCAGGTATCTTCGAGCCAGCCACGATCGACATTGATCTGGCTATCTTGCTGACAAAAG
CAAGAGAACATAGCGTTGCCTTGGTAGGTCCAGCGGCGGAGGAACTCTTTGATCCGGTTCCTGAACAGGAT
CTATTTGAGGCGCTAAATGAAACCTTAACGCTATGGAACTCGCCGCCCGACTGGGCTGGCGATGAGCGAAA
TGTAGTGCTTACGTTGTCCCGCATTTGGTACAGCGCAGTAACCGGCAAAATCGCGCCGAAGGATGTCGCTG
CCGACTGGGCAATGGAGCGCCTGCCGGCCCAGTATCAGCCCGTCATACTTGAAGCTAGACAGGCTTATCTT
GGACAAGAAGAAGATCGCTTGGCCTCGCGCGCAGATCAGTTGGAAGAATTTGTCCACTACGTGAAAGGCGA
GATCACCAAGGTAGTGGGCAAAGAACAAAAACTCATTTCTGAAGAAGACTTGTAACTGCAGAGACATTAGCA
GATAAATTAGCAGGAAATAAAGAAGGATAAGGAGAAAGAACTCAAGTAATTATCCTTCGTTCTCTTAATTGAAT
TGCAATTAAACTCGGCCCAATCTTTTACTAAAAGGATTGAGCCGAATACAACAAAGATTCTATTGCATATATTTT
GACTAAGTATATACTTACCTAGATATACAAGATTTGAAATACAAAATCTAACATACACCTTGGTTGACACGAGTA
TATAAGTCATGTTATACTGTTGAATAACAAGCCTTCCATTTTCTATTTTGATTTGTAGAAAACTAGTCTGCTTCC
GAATCCGTGATGATTAAATAAACCAAGATTTTACCATGACTGCAATTTTAGAGAGACGCGAAAGCGAAAGCCT
ATGGGGTCGCTTCTGTAACTGGATAACTGCTAGCAGTAAAGGAGAAGAACTTTTCACTGGAGTTGTCCCAAT
TCTTGTTGAATTAGATGGTGATGTTAATGGGCACAAATTTTCTGTCAGTGGAGAGGGTGAAGGTGATGCAAC
ATACGGAAAACTTACCCTTAAATTTATTTGCACTACTGGAAAACTACCTGTTCCTTGGCCAACACTTGTCACTA
CTTTCTCTTATGGTGTTCAATGCTTTTCAAGATACCCAGATCATATGAAGCGGCACGACTTCTTCAAGAGCGC
CATGCCTGAGGGATACGTGCAGGAGAGGACCATCTCTTTCAAGGACGACGGGAACTACAAGACACGTGCT
GAAGTCAAGTTTGAGGGAGACACCCTCGTCAACAGGATCGAGCTTAAGGGAATCGATTTCAAGGAGGACGG
AAACATCCTCGGCCACAAGTTGGAATACAACTACAACTCCCACAACGTATACATCACGGCAGACAAACAAAA
GAATGGAATCAAAGCTAACTTCAAAATTAGACACAACATTGAAGATGGAAGCGTTCAACTAGCAGACCATTAT
CAACAAAATACTCCAATTGGCGATGGCCCTGTCCTTTTACCAGACAACCATTACCTGTCCACACAATCTGCCC
TTTCGAAAGATCCCAACGAAAAGAGAGACCACATGGTCCTTCTTGAGTTTGTAACAGCTGCTGGGATTACAC
ATGGCATGGATGAACTATACAAATAAGCTCTAGCTAGAGCGATCCTGGCCTAGTCTATAGGAGGTTTTGAAAA
GAAAGGAGCAATAATCATTTTCTTGTTCTATCAAGAGGGTGCTATTGCTCCTTTCTTTTTTTCTTTTTATTTATTT
ACTAGTATTTTACTTACATAGACTTTTTTGTTTACATTATAGAAAAAGAAGGAGAGGTTATTTTCTTGCATTTATT
CATGGGGGATCAAAGCTTGATACTTTAACTGCCCCTATCGGAAATAGGATTGACTACCGATTCCGAAGGAACT
GGAGTTACATCTCTTTTCCATTCAAGAGTTCTTATGCGTTTCCACGCCCCTTTGAGACCCCGAAAAATGGACA
AATTCCTTTTCTTAGGAACACATACAAGATTCGTCACTACAAAAAGGATAATGGTAACCCTACCATTAACTACTT
CATTTATGAATTTCATAGTAATAGAAATACATGTCCTACCGAGACAGAATTTGGAACTTGCTATCCTCTTGCCTA
GCAGGCAAAGATTTACCTCCGTGGAAAGGATGATTCATTCGGATCGACATGAGAGTCCAACTACATTGCCAG
AATCCATGTTGTATATTTGAAAGAGGTTGACCTCCTTGCTTCTCTCATGGTACACTCCTCTTCCCGCCGAGCC
CCTTTTCTCCTCGGTCCACAGAGACAAAATGTAGGACTGGTGCCAACAATTCATCAGACTCACTAAGTCGGG
ATCACTAACTAATACTAATCTAATATAATAGTCTAATATATCTAATATAATAGAAAATACTAATATAATAGAAAAGAAC
TGTCTTTTCTGTATACTTTCCCCGGTTCCGTTGCTACCGCGGGCTTTACGCAATCGATCGGATTAGATAGATAT
CCCTTCAACATAGGTCATCGAAAGGATCTCGGAGACCCACCAAAGTACGAAAGCCAGGATCTTTCAGAAAAC
GGATTCCTATTCAAAGAGTGCATAACCGCATGGATAAGCTCACACTAACCCGTCAATTTGGGATCCAAATTCG
AGATTTTCCTTGGGAGGTATCGGGAAGGATTTGGAATGGAATAATATCGATTCATACAGAAGAAAAGGTTCTC
TATTGATTCAAACACTGTACCTAACCTATGGGATAGGGATCGAGGAAGGGGAAAAACCGAAGATTTCACATGG
TACTTTTATCAATCTGATTTATTTCGTACCTTTCGTTCAATGAGAAAATGGGTCAAATTCTACAGGATCAAACCT
ATGGGACTTAAGGAATGATATAAAAAAAAGAGAGGGAAAATATTCATATTAAATAAATATGAAGTAGAAGAACCC
AGATTCCAAATGAACAAATTCAAACTTGAAAAGGATCTTCCTTATTCTTGAAGAATGAGGGGCAAAGGGATTG
ATCAAGAAAGATC

### (B) VIGS constructs

Sequence of Nt HCF244 cDNA fragment for VIGS

GCTATGTGGCTTCATGCAGGGCCTTATCGGTCAATATGCAGTGCCTATATTAGAAGAGAAATCTGTATGG
GGAACTGATGCTCCCACTCGAATAGCATACATGGACACCCAAGATATTGCTCGCTTGACATTCATAGCTT

TACGCAATGAGAATATTAATGGGAAGCTTCTCACTTTTGCTGGGCCTCGGGCATGGACAACCCAAGAGGT
GATAACATTGTGCGAGAGACTTGCCGGCCAAGATGCCAATGTGACTACAGTGCCTGTCTCAGTTTTGCGA
TTGACACGGCAGCTGACTCGGTTGTTTGAGTGGACGAATGATGTTGCTGATAGATTGGCATTTTCAGAGG
TTTTGACAAGCGATACTGTCTTCTCAGTTCCTATGACTGAGACATATAGCCTTCTCGGTGTGGATGCAAA
AGACGTCAGTTCACTGGAGAAGTATCTGCAGGATTATTTCACCAACATACTGAAGAAGTTGAAAGACCTT
AAGGCTCAATCAAAGCAAACAGATATTTTCTTTTGAGAGGAATACTTGTTTATGTTGTTGTAAAAGTAAT
ATACACTGTGACAGGGATACATCAGCAGAGGCATGCTCCT

Sequence of Nt HCF173 cDNA fragment for VIGS

CGTGGAGTTGGTGACAGGGGACGTTGGTGATCCTTCTAGTCTAAAAGATGCAGTGCAAGGCTGCAGCAAA
ATCATCTATTGTGCCACTGCTCGTTCTTCAATCACGGGTGATCTCATCAGAGTTGATCATCAAGGGGTTT
ACAATCTAACCAAAGCTTTGCAGGACTACAATAATAAACTAGCACAGCAACGCGCCGGGAAGAGTAGCAA
AAGCAAGCTTTTAATTGCAAAATTCAAGTCTGAAGATTCATTGAACGGGTGGGAAGTCCGTCAAGGGACA
TATTTCCAGGATGTGGTTGCTTCTAAGTATGATGGAGGAATGGATGCCACGTTTGAGTTTACTGAAAGCG
GAGAGACTATTTTTTCAGGATATGTTTTCACAAGAGGAGGCTATGTTGAATTGTCAAGAAAACTGTCACT
TCCTTTAGGTTATACTCTTGACAGGTACGAGGGTCTAGTATTCTCTGTTGGTGGGAATGGGAGATCTTAT
GTTGTAATTCTTGAAGCTGGTCCTTCAGCAGATACAACCCAAAGCAAACTGTATTTTGCCAGAATTAGCA
CAAAAGCAGGATTTTGCAGGGTGAGAGTTCCATTTTCCTC

**(C) Sequences encoding Arabidopsis HCF173-HA and HCF173Trunc-HA.**

Nucleotides deleted in *HCF173Trunc-HA* are in bold and the sequence encoding the HA tag is highlighted in yellow.

ATGGTTGGCTCAATTGTGGGTTCCAACATGGCTGCCACTGACGCTAGATTCCTTTCAAGTAACTTCGGGAATAGCTTTAGCA
TTAATACTCGAATACATCGATTCCATGACAGATCTCAGATTGTGATTCCCCGTGCTCAGAGTTCTAGCTCTCCTAGTCCATC
CCCCCCGTCCGATAAAAAGAAGACAAAAACTAGGCCAGGTACTATAACAACCAAAGAATCAGAAGAGACCGTCGCGAAAAAG
CTGGATGTAGCCCCACCGTCACCCCAGTCTCCGCCAAGTCCCCCTACACTAAAGCTAGATGACGTGAACCCGGTTGGACTTG
GTAGACGATCACGTCAGATTTTTGACGAGGTGTGGCGTAAGTTTAGCGGACTGGGGCAGATGTCAAGAACGACTAGACCGGA
TGAGCAGGAAACACTCGACAGCTTACTGATAAGAGAAGGTCCGATGTGTGAATTCGCGGTTCCGGGTGCACAGAACGTGACG
GTTCTTGTAGTTGGTGCTACGTCCAGAATCGGCAGGATTGTGGTACGAAAGCTAATGCTCCGTGGATACACCGTCAAGGCCC
TAGTTAGGAAACAGGATGAGGAGGTCATGTCCATGTTGCCACGATCTGTAGATATTGTAGTAGGCGATGTTGGCGAACCCTC
TACGTTGAAGTCAGCTGTCGAAAGTTGCAGCAAGATTATTTATTGTGCGACCGCTAGAAGCACGATTACTGCAGATCTCACG
CGAGTCGATCACCTAGGAGTCTACAATCTGACAAAGGCCTTTCAAGATTACAATAACAGGTTGGCCCAGCTCCGTGCTGGCA
AGAGTTCTAAAAGCAAACTATTGCTGGCAAAGTTTAAGAGTGCCGAGAGCCTTGACGGGTGGGAAATAAGACAAGGTACGTA
CTTTCAAGACACTACCGCTAGTAAATATGATGGGGGCATGGACGCAAAATTCGAGTTCACGGAGACCGAGAGGGCTGAGTTT
TCCGGATATGTCTTTACGAGAGGCGGCTACGTTGAGCTGTCTAAAAAACTATCCCTGCCGCTAGGAACGACTTTGGACAGAT
ACGAGGGTCTCGTACTGTCTGTAGGCGGAAATGGACGAAGCTATGTCGTGATACTGGAAGCGGGTCCCTCTTCCGATATGTC
ACAGAGTAAACAATACTTCGCACGAATCTCTACTAAAGCGGGTTTCTGCAGAGTGCGAGTACCGTTTAGTGCATTTCGTCCT
GTGAACCCTGAGGACCCCCCGCTTGATCCTTTTCTCGTTCATACTCTTACTATAAGATTTGAGCCGAAGCGACAAAGGCCTG
TCGATGGCTTAGCTGGTGCCCAGCAGGATCTCAGGAGCTTTTCACTCGTCTTTGAATACATCAAGGCATTGCCGGCTGGTCA
GGAGACGGATTTTATCCTAGTAAGCTGTACAGGATCTGGGGTGGAAGCGAACAGGAGAGAACAAGTTCTCAAAGCTAAACGA
GCAGGCGAGGATTCCCTGCGAAGATCTGGCCTGGGCTATACTATCATTAGACCTGGACCTCTGAAGGAGGAACCTGGAGGAC
AACGTGCACTAATCTTCGATCAGGGCAACAGGATTTCTCAGGGTATTAGCTGCGCCGATGTTGCGGATATTTGCGTTAAGGC
GTTGCATGATAGCACCGCGAGGAATAAATCTTTCGACGTTTGCCACGAATACGTAGCGGAACAGGGAATCGAGTTATATGAA
CTC**GTCGCTCATTTGCCAGATAAGGCAAACAACTACTTGACACCCGCACTGAGTGTCCTGGAGAAGAACACA**GGTACCTATC
CATATGACGTCCCTGACTACGCGTACCCGTACGATGTCCCCGATTATGCGTATCCTTACGATGTCCCGGACTATGCTTAA

**Supplementary Figure S2. Characteristics of transplastomic plants**
(A) Visible phenotypes of transplastomic *psbA-gfp, psbA$^{mut-}$gfp* and *psbA-pagA* plants. GFP fluorescence was imaged by illumination with UV light (right).
(B) Southern blot hybridizations demonstrating homoplasmy of the *psbA-gfp, psbA$^{mut-}$gfp* lines. The blot was probed with an oligonucleotide complementary to plastid 16S rRNA.
(C) RNA gel blot hybridizations demonstrating loss of *gfp* mRNA in independent *psbA$^{mut-}$gfp* transformed lines. The blot was hybridized with a probe that is complementary to the first 50 nt of the *psbA* ORF, which is found in both the *psbA* and *gfp* transcripts. The same blot was stained with methylene blue to image rRNAs as a loading control.

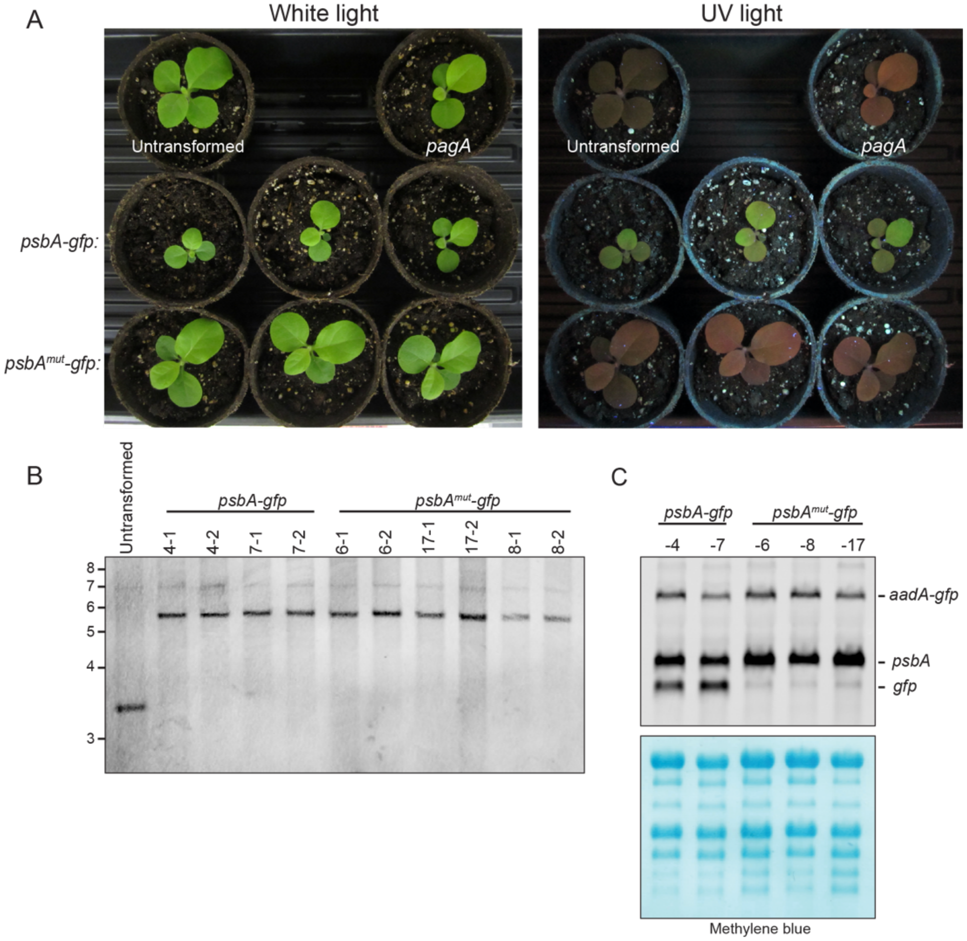

**Supplementary Figure S3. Data in support of polysome and ribo-seq analyses**

(A) Replicate polysome analysis of *psbA-gfp* line. Leaf tissue was harvested after one hour of dark adaptation or after 15 minutes of reillumination. This experiment is a replicate of that shown in Figure 2A but used tissue from an independent transformed line. GFP and *psbA* mRNAs were detected with probes complementary to their respective ORFs.

(B) Replicate polysome analysis of the *pagA* line. Leaf tissue was harvested after one hour of dark adaptation or after 15 minutes of reillumination. This experiment is a replicate of that shown in Figure 2B. *pagA* and *psbA* mRNAs were both detected with a probe complementary to the *psbA* 5' UTR.

(C) Replicate ribo-seq analyses of *psbA-gfp* plants. Plants were grown for 6-weeks and a mature leaf from different plants at a matched developmental stage was harvested at midday, after 1 hour in the dark or after 15 min of reillumination. Values represent the mean of two biological replicates ± SEM.

(D) qRT-PCR analysis demonstrating similar reporter mRNA concentrations in lysates from dark-adapted and illuminated ribo-seq samples used in Figures 4D and 5D. The mean ± SD of four technical replicates of each of two biological replicates is presented.

(E) RNA gel blot hybridization illustrating *psbA* mRNA abundance in samples used for ribo-seq analysis of dark-adapted (D) and reilluminated (R) Arabidopsis HCF173-HA and HCF173Trunc-HA transgenic lines. Supports Figure 8D.

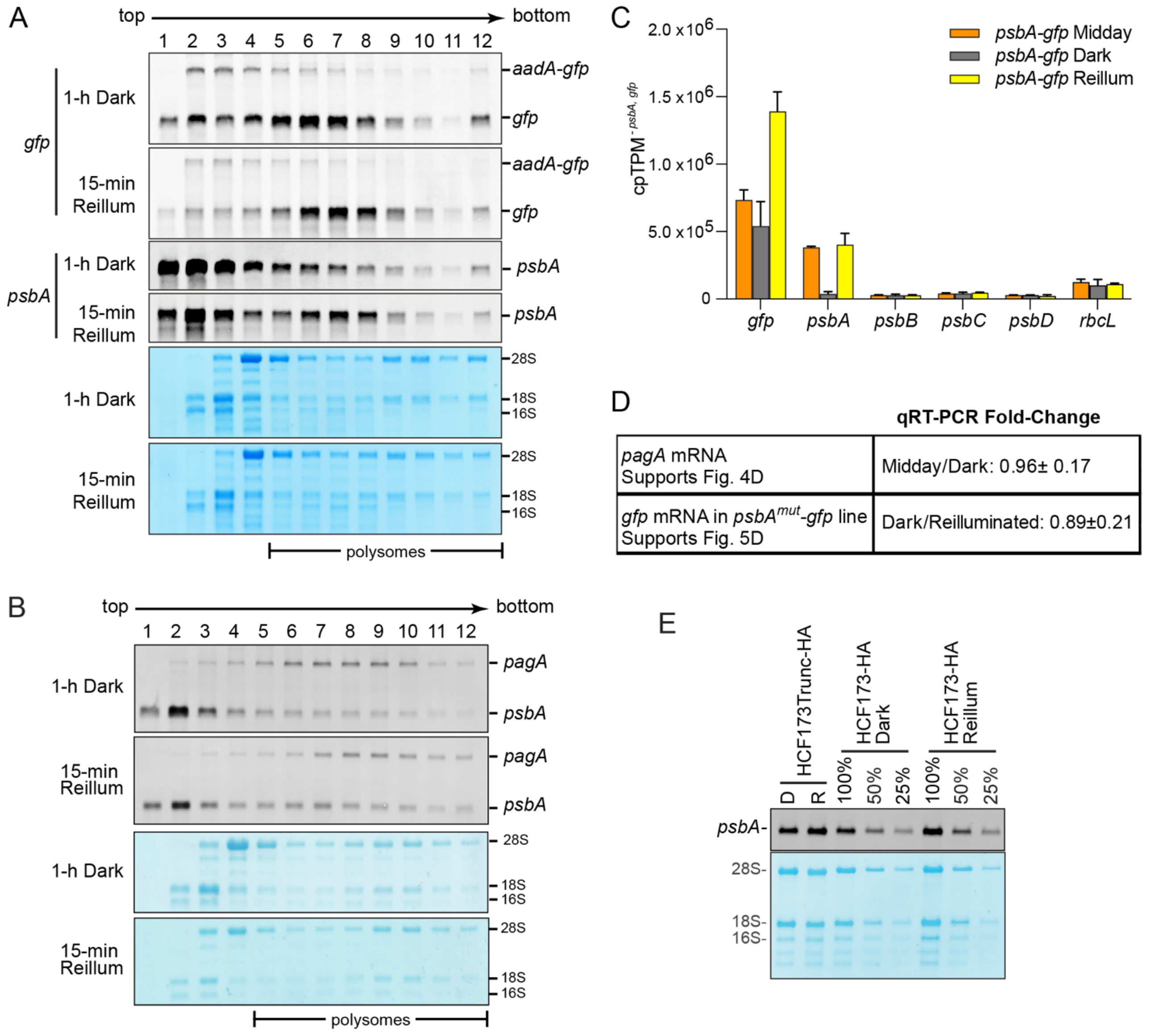

**Supplementary Figure S4. Abundance of HCF244 and HCF173 during short-term light shifts.** Maize (A) and Arabidopsis (B) seedling leaf tissue was harvested at midday, after 1-h in the dark, and following 15-min of reillumination. Total lysates were resolved by SDS-PAGE, transferred to nitrocellulose, and probed with the indicated antibodies. The same blots were stained with Ponceau S to illustrate relative sample loading.

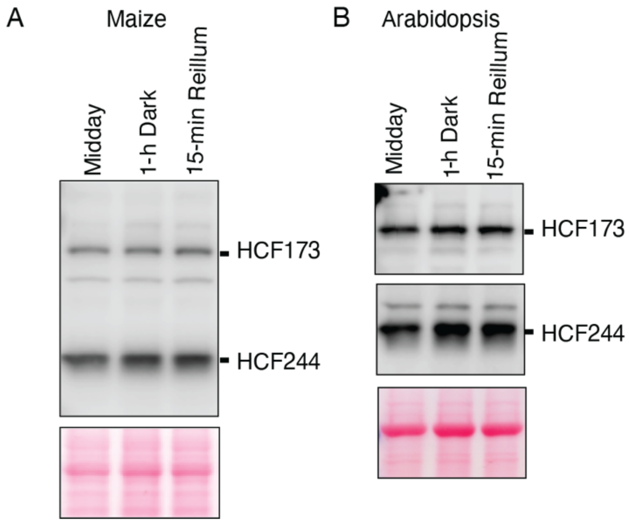

**Supplementary Figure S5. Multiple sequence alignment of HCF173 orthologs and paralogs.** (A) Arabidopsis (At, AT1G16720.1), maize (Zm, GRMZM2G397247_P03) and Chlamydomonas (Cr, Cre13.g578650_4532.1) HCF173 orthologs are aligned with the HCF173 paralogs in Arabidopsis (AT4G18810.1) and maize (Zm00001eb204900_T002). Green and cyan mark the two segments of the SDR domain, magenta marks the CIA30 domain, and orange marks the carboxy terminal tail conserved specifically in HCF173 orthologs. The carboxy terminus of the HCF173Trunc mutant is marked with a delta. (B) Helical wheel projection of HCF173's carboxy-terminal tail rendered with NetWheels (https://neutrophil-proteome.shinyapps.io/netwheels/).

A

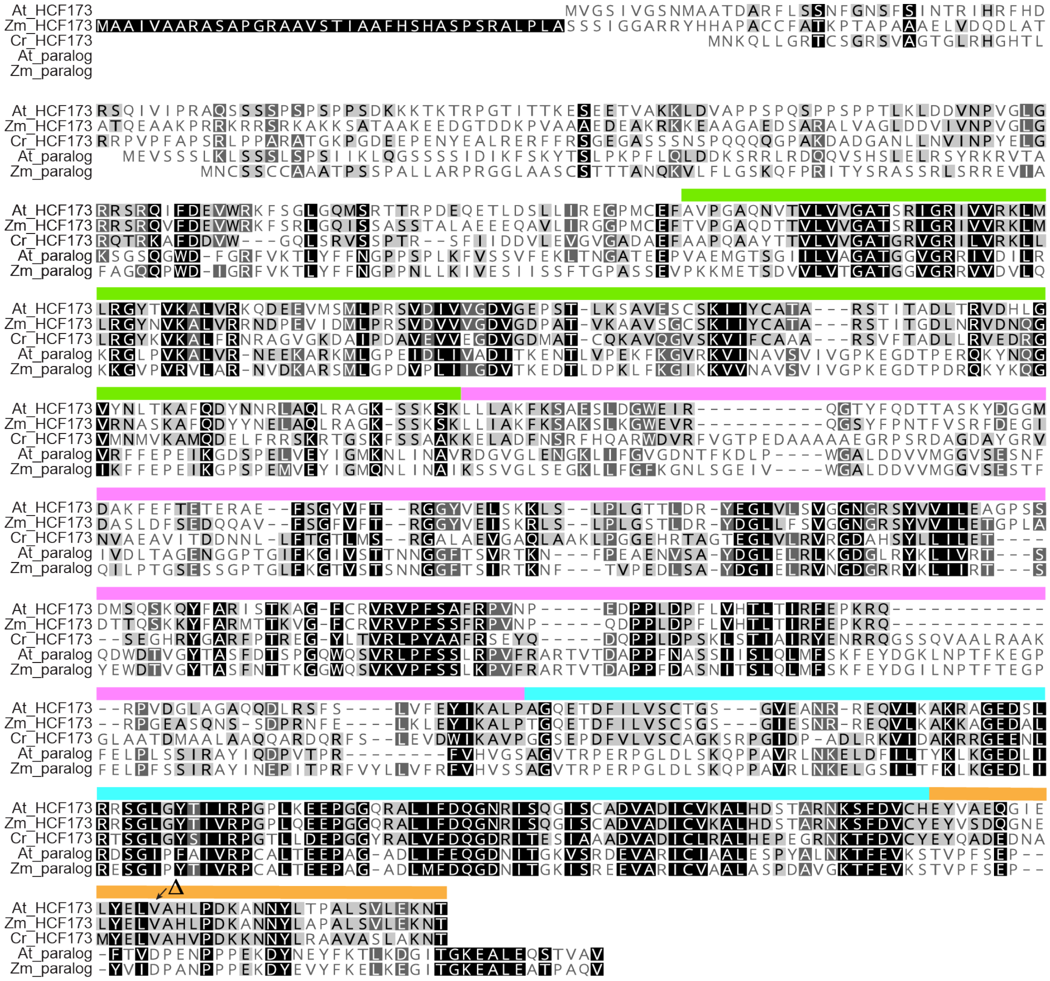


B

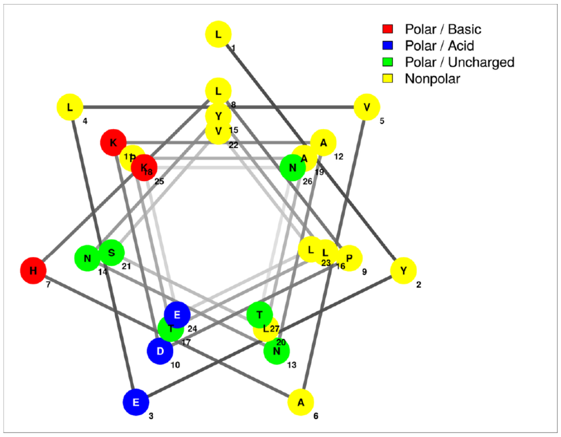
